# Transcriptomic profiling of ‘*Candidatus* Liberibacter asiaticus’ in different citrus tissues reveals novel insights into Huanglongbing pathogenesis

**DOI:** 10.1101/2024.09.02.610751

**Authors:** Amelia H. Lovelace, Chunxia Wang, Amit Levy, Wenbo Ma

## Abstract

‘*Candidatus* Liberibacter asiaticus’ (Las) is a gram-negative bacterial pathogen associated with citrus huanglongbing (HLB) or greening disease. Las is transmitted by the Asian citrus psyllid (ACP) where it colonizes the phloem tissue, resulting in substantial economic losses to citrus industry worldwide. Despite extensive efforts, effective management strategies against HLB remain elusive, necessitating a deeper understanding of the pathogen’ s biology. Las undergoes cell-to-cell movement through phloem flow and colonizes different tissues in which Las may have varying interactions with the host. Here, we investigate the transcriptomic landscape of Las in citrus seed coat vasculatures, enabling a complete gene expression profiling of Las genome and revealing unique transcriptomic patterns compared to previous studies using midrib tissues. Comparative transcriptomics between seed coat, midrib and ACP identified specific responses and metabolic states of Las in different host tissue. Two Las virulence factors that exhibit higher expression in seed coat can suppress callose deposition. Therefore, they may contribute to unclogging sieve plate pores during Las colonization in seed coat vasculature. Furthermore, analysis of regulatory elements uncovers a potential role of LuxR-type transcription factors in regulating Liberibacter effector gene expression during plant colonization. Together, this work provides novel insights into the pathogenesis of the devastating citrus HLB.

**Funding:** This work is supported by USDA National Institute of Food and Agriculture award No. 2020-70029-33197 to W.M and A.L.

## INTRODUCTION

Citrus Huanglongbing (HLB), also known as citrus greening disease, is the most devastating citrus disease in the world, resulting in billions of dollars in economic losses annually (Wang 2019, Singerman and Rogers 2020). This disease is primarily caused by gram-negative bacteria belonging to ‘*Candidatus* Liberibacter’ species, including *Ca.* Liberibacter asiaticus (Las), *Ca.* Liberibacter americanus (Lam), and *Ca.* Liberibacter africanus (Laf), with Las being the most widespread and impactful to the citrus industry (Bové 2006). *Ca.* Liberibacter species are specialized, obligate, phloem-colonizing bacteria transmitted by psyllids as the insect vector. More specifically, Las is vectored by the Asian citrus psyllid (ACP, *Diaphorina citri*) (Wang 2017).

HLB is considered a pathogen-triggered immune disease in that Liberibacter colonization of the phloem sieve elements stimulates systemic immune responses including callose deposition and production of reactive oxygen species (ROS) (Ma et al. 2022, Pitino et al. 2017). This response leads to phloem degeneration and blockage of phloem sap movement from source to sink tissues (Schneider 1968), resulting in HLB symptoms such as leaf mottling and chlorosis, stunted growth, deformed fruit, and premature fruit drop (Gabriel et al. 2020). Despite the enormous efforts that have been invested to controlling HLB, no curative treatment or management strategies have been successfully implemented to eliminate Liberibacter from infected citrus trees (Barnett et al. 2019, Huang et al. 2021). A deeper understanding of the basic biology of this bacterial pathogen is urgently needed to develop sustainable methods so that global citrus industry can be protected from HLB.

Recent studies have discovered differences in Las colonization in different parts of citrus tree. Cell-to-cell movement of Las was observed in the phloem, potentially via phloem flow, and Las was present at a higher titer in some sink tissues compared to source tissues (Achor et al. 2020). More specifically, the seed coat tissue, which is enriched for vasculature and functions to deliver nutrients to the developing embryo, was found to have a high accumulation of Las cells. Interestingly, sieve plates pores were not occluded in infected seed coat tissue, correlated with the reduced callose or P-protein production/deposition. In contrast, callose deposition leads to blockage of phloem sieve plate pores in the midrib tissues of infected leaves, suggesting that Las may manipulate plant immune responses in a tissue-specific manner. Consistent with this possibility, infected seed coats had a reduced ROS accumulation after Las infection while ROS was induced in infected leaves (Bernardini et al. 2022).

Virulence factors are molecules produced by a pathogen that contribute to their ability to infect host and cause disease. Genome sequences of several Las isolates have allowed for the mining of virulence proteins (Thapa et al. 2020). As *Ca.* Liberibacter species do not encode the type III secretion system, virulence proteins may be secreted into the phloem through the general Sec secretion system. A core set of Sec-dependent effectors (SDEs) have been predicted from Las isolates (Pagliaccia et al. 2017, Thapa et al. 2020) of which the virulence activity has been characterized for a few, namely SDE1, SDE3, SDE15, and most recently CLIBASIA_03230 (Clark et al. 2018, Clark et al. 2020, Shi et al. 2023, Pang et al. 2020, Zhang et al. 2024). In addition to SDEs, other virulence factors have also been demonstrated to play roles in manipulating host defenses. For example, two 1-Cys peroxiredoxins (*prx5* and *bcp*) and a prophage-encoded peroxidase (SC2_gp095) in Las contribute to increased bacterial fitness via ROS detoxification (Jain et al. 2015, Jain et al. 2018). Additionally, a salicylic acid (SA) hydroxylase was shown to suppress SA accumulation and pathogenesis-related gene expression in citrus (Li et al. 2017).

While several virulence factors have been characterized, a general understanding of Las pathogenesis is lacking. Transcriptome analysis is a powerful tool for providing insight into the basic biology of a pathogen during host colonization. However, this has been challenging for *Ca.* Liberibacter due to their overall low titer and uneven distribution in the citrus phloem tissue. Previously, we employed a bacterial cell enrichment method using infected leaf midrib tissues, which allowed us to detect the expression of up to 69% of the total Las genes (De Francesco et al. 2022). Comparative analysis of this dataset with the Las transcriptome from the ACP enabled the establishment of specific processes important for colonizing each host. However, there was still a significant portion of Las genes that were undetectable for their expression within citrus tissue. Furthermore, how Las may adapt to different citrus niches, such as in leaf midrib vs seed coat tissues, was unknown.

Here, we determined the Las transcriptome in citrus seed coat tissues, which harbored high Las titers. This analysis resulted in the detection of 99% of the entire Las-encoded genes and a complete coverage of Las gene expression profile in planta. We compared differentially expressed genes (DEGs) of Las when colonized different citrus tissues, identifying pathways and cellular processes that were distinctly enriched for the colonization in leaf midrib vs seed coat. Additionally, we identified and characterized key virulence factors that may play a role in manipulating host defenses in a tissue-specific manner. The full coverage of Las gene expression profile also allowed us to identify a LuxR motif, which was enriched in *Ca.* Liberibacter effector promoters. A similar motif is also enriched in the effector gene promoters in Lso, suggesting the potential role of LuxR-type transcription factors in regulating *Ca.* Liberibacter virulence in planta. This study advances a basic understanding of how Las may adapt to different host environments and manipulate host defenses and revealed regulatory components that may contribute to transcription reprogramming.

## RESULTS

### Determination of Las transcriptomics using seed coat tissues

Our understanding of Las biology during phloem colonization is limited by the current transcriptomic data, which does not provide a full coverage of the genome. We sought to overcome this limitation by investigating the transcriptional landscape of Las using isolated vasculature from infected citrus seed coats, which are enriched for Las. Las infection was confirmed in seed coat tissues and their titer was determined by reverse transcription quantitative PCR (RT-qPCR) using *16S* rRNA as primers (Table 1, Supplemental Table 2). The cycle threshold (Ct) values were compared to that of healthy tissue, which had values of >30. Relative abundance of Las was calculated as ΔCt (= Ct_Las 16S_ -Ct_Citrus Fbox_) using the Ct value of a citrus housekeeping gene (encoding an F box protein) for normalization. The results show a much higher Las titers in the seed coat tissues compared to infected leaf midrib samples analyzed in our previous study (Table 1).

**Table 1.** *Candidatus* Liberibacter asiaticus (Las) titer and relative abundance in different Grapefruit tissues.

| Sample<sup>a</sup> | Las titer<sup>b</sup> | Healthy/Diseased | Relative abundance of Las<sup>c</sup> |
| --- | --- | --- | --- |
| midrib_1* | 20.67 | Diseased | -6.41 |
| midrib_2* | 22.44 | Diseased | -3.24 |
| midrib_3* | 22.67 | Diseased | -2.28 |
| seedcoat_1 | 13.35 | Diseased | -11.68 |
| seedcoat_2 | 13.48 | Diseased | -11.63 |
| seedcoat_3 | 14.82 | Diseased | -10.39 |
| seedcoat_4 | 13.54 | Diseased | -3.52 |
| seedcoat_5 | 31.20 | Healthy | 4.92 |
| seedcoat_6 | 30.53 | Healthy | 5.37 |
| seedcoat_7 | 31.19 | Healthy | 5.94 |

<sup>a</sup> Grapefruit tissue type and biological replicate for huanglongbing-infected or healthy samples. \*Previously published in De Francesco et al. 2022.
<sup>b</sup> Cycle threshold (Ct) values of Las 16S rRNA determined by reverse transcription quantitative PCR.
<sup>c</sup> Relative abundance of Las to citrus genes was calculated as $\Delta Ct = (Ct_{\text{Las 16S}} - Ct_{\text{Citrus Fbox}})$ . Negative values indicate higher relative abundance of Las.

We then subjected the seed coat samples to RNA-seq. Compared to the midrib, RNA extracted from the seed coat samples detected a higher number of Las transcripts with 5.1- and 4.9-fold increase in average read depth and genome coverage, respectively (Table 2). Additionally, almost all the Las genes were detected in seed coat RNA samples (up to 99.10%) compared to up to 68.68% coverage in the midrib samples (Table 2). Therefore, the seed coat samples allowed a near complete detection of all genes encoded in the Las genome.

**Table 2.** RNA sequencing results from Las infected samples.

| Sample<sup>a</sup> | Read Depth<sup>b</sup><br>(x1,000) | Mapped reads<sup>c</sup><br>(%) | Coverage<sup>d</sup> | Las genes detected<sup>e</sup><br>(%) |
| --- | --- | --- | --- | --- |
| seedcoat_1 | 70.2 | 0.164 | 8.58 | 98.11 |
| seedcoat_2 | 139.7 | 0.330 | 17.07 | 99.10 |
| seedcoat_3 | 65.0 | 0.153 | 7.95 | 96.40 |
| seedcoat_4 | 33.3 | 0.028 | 2.71 | 93.07 |
| midrib_1.1* | 15.9 | 0.011 | 1.95 | 62.38 |
| midrib_1.2* | 16.2 | 0.011 | 1.98 | 68.68 |
| midrib_2.1* | 17.3 | 0.015 | 2.11 | 62.38 |
| midrib_2.2* | 17.5 | 0.015 | 2.14 | 31.86 |
| midrib_3.1* | 11.9 | 0.011 | 1.46 | 43.29 |
| midrib_3.2* | 11.7 | 0.011 | 1.43 | 44.55 |
| ACP_1* | 41.8 | 0.033 | 5.11 | 95.77 |
| ACP_2* | 60.8 | 0.047 | 7.43 | 96.40 |
| ACP_3* | 423.3 | 0.214 | 51.73 | 99.64 |
| ACP_4* | 2216.4 | 1.373 | 270.88 | 100.00 |

<sup>a</sup> Grapefruit tissue type, biological and technical replicate for huanglongbing-infected samples. ACP, Asian citrus psyllid. \*Previously published in De Francesco et al. 2022.
<sup>b</sup> Read depth calculated as total reads mapped to *Candidatus Liberibacter asiaticus* (Las) psy62 genome (NC\_012985.3) from HISAT2 v.2.1.0 output.
<sup>c</sup> Percent of total sequenced read that mapped to the Las psy62 genome (NC\_012985.3) from HISAT2 v.2.1.0 output.
<sup>d</sup> Coverage was calculated as the product of read depth and read length divided by the total genome size, in which read length is 150 bp for all midrib samples, all ACP samples, and seedcoat samples 1-3 and 100 bp for seedcoat\_4 sample. The genome size for Las psy62 is 1,227,328 bp.
<sup>e</sup> Percentage of Las psy62 genes with mapped reads out of the total number of genes in the genomes (1,111 genes).

### Differential expression of Las genes in citrus midribs, seed coats, and psyllids

Principle component analysis (PCA) of gene expression profiles represented as the normalized gene counts for all samples was performed for quality assessment and an overall comparison between sample types (Supplemental Dataset 1). Transcriptome profiles of Las from the same tissue/host type were nicely grouped together whereas those from different tissue/host types were clearly separated (Figure 1A), indicating distinct transcriptomic signatures of Las in the tissues, especially in seed coat vs leaf midrib.

**Figure 1.**
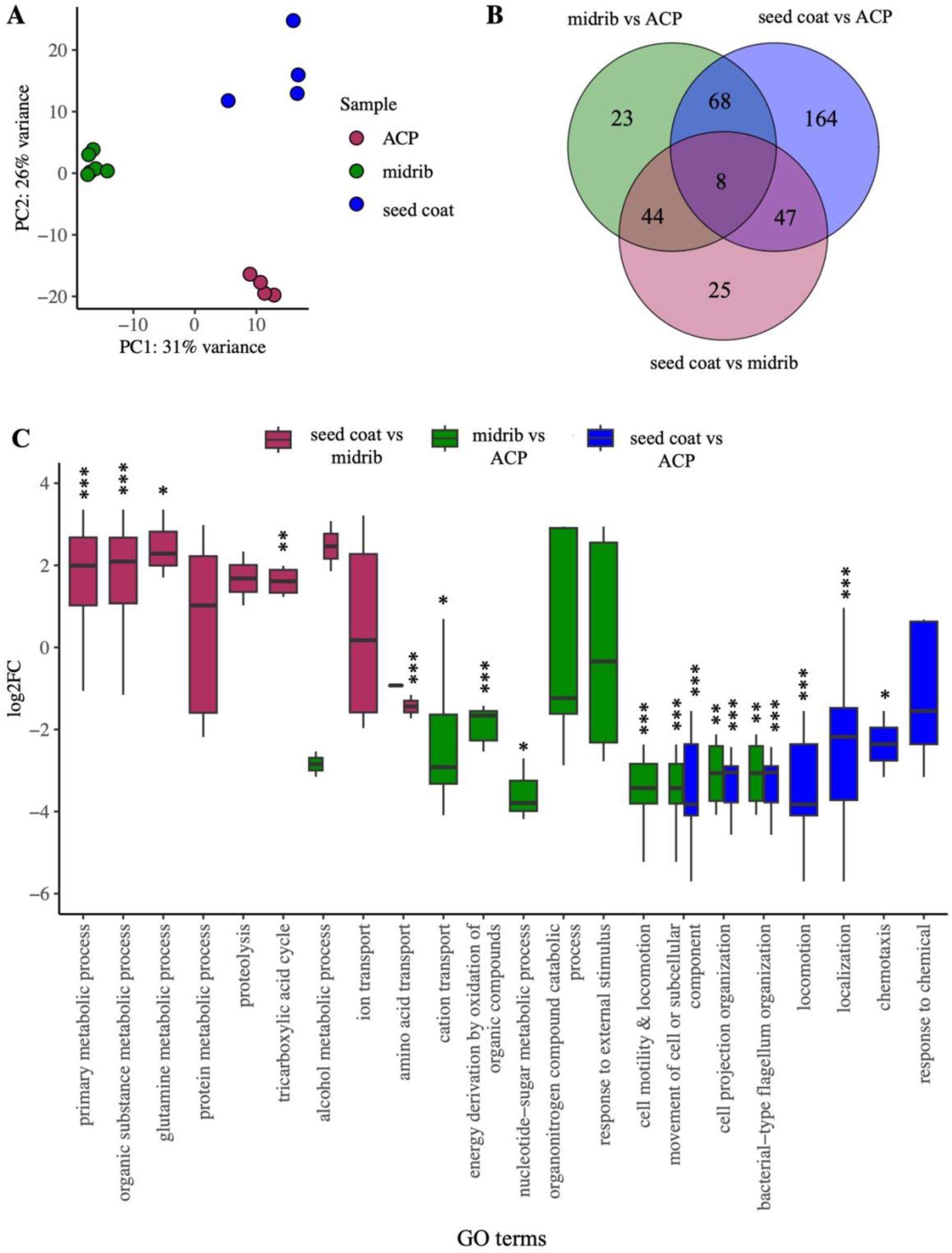
Differentially expressed genes (DEGs) of *Candidatus* Liberibacter asiaticus (Las) when colonizing citrus midribs, citrus seed coats, and Asian citrus psyllid (ACP). **A.** Principal component analysis of Las gene expression profiles measured in log-transformed gene counts in different host tissues including citrus midribs (green), citrus seed coats (blue), and ACP (maroon). Gene counts from RNA sequencing data was transformed using the rlog function in DESeq v.1.30.1. **B.** Venn diagram of shared DEGS of Las using pairwise comparisons. DEGs were identified as genes with |log_2_ fold change| (log_2_FC) > 0 and an adjusted p-value < 0.05. For a full DEG list, see Supplemental Dataset S2. **C.** Significantly enriched Gene Ontology (GO) terms found in DEGs using pairwise comparisons. Significantly enriched genes for each GO terms are plotted based on their log_2_FC in citrus midrib compared to ACP (green), citrus seed coat compared to ACP (blue), and midrib compared to seed coat (maroon). For a full GO list, see Supplemental Dataset S3a-c. Asterisks indicate log_2_FC values significantly different from 0 (two-tailed *t* test at * p < 0.05, ** p < 0.01, *** p < 0.001).

Pair-wise comparisons were made between the transcriptomes of Las in citrus (midribs and seed coats) and in ACP as well as between the two citrus tissue types to identify genes that may contribute to bacterial colonization in these different host environments. For each comparison, we determined differentially expressed genes (DEGs), which are defined as genes with significant differences in relative expression between the sample types (|log2FC| > 0 and adjusted *P* < 0.05) (Figure 1B, Supplemental Dataset 2). The midrib and seed coat samples had 143 and 287 total DEGs when compared to ACP samples respectively, with 76 DEGs shared in both samples, indicating their contribution to in planta colonization regardless of tissue types.

We also detected unique DEGs in midrib or seed coat samples through pairwise comparisons with ACP. In midrib samples, 23 genes were differentially expressed compared to ACP while 164 were identified in the seed coat samples. Ninety-one genes were specifically upregulated in seed coats. These include genes encoding virulence factors such as putative SDEs CLIBASIA_03230, CLIBASIA_04560, and CLIBASIA_04580, as well as a peroxiredoxin, Bcp. In addition, 73 genes were specifically downregulated, such as SDEs CLIBASIA_00520 and CLIBASIA_00525, components of the Sec machinery SecE and SppA, and genes involved in NADH-quinone oxidoreductase respiration and ribosome biosynthesis. These results reveal changes in Las cellular processes that were undetectable in the previous analysis using midrib sample, in part due to the overall low abundance of Las transcripts in the RNA-seq dataset. However, we also detected biological processes downregulated in seed coats that are not due to fewer detectable genes in the midrib samples. In fact, 180 genes were undetectable in any midrib samples, of which only 17 were found to be DEGs in seed coats. The disparity in unique DEGs between the two citrus tissue types when individually compared to ACP indicates tissue-specific interactions of Las in citrus.

Pairwise comparison between the two citrus tissue-types identified 124 DEGs of which 25 were not found when comparing citrus vs ACP (Figure 1B). Seventeen of the 25 unique DEGs were upregulated. These genes are involved in transcription and the TCA cycle, suggesting that Las induces shifts in metabolism and reprogramming at the RNA level during colonization of the seed coats. Despite the small number of genes identified, these results indicate tissue-specific changes in Las gene expression to adapt to different phloem niches.

We then subjected DEGs from each comparison to GO analysis to better understand enriched cellular processes in the different host environment (Figure 1C, Supplemental Dataset 3). The significant DEGs found within each enriched GO term were also used to determine if the GO terms were significantly upregulated or downregulated based on their log_2_FC values. In seed coat tissues, genes related to primary, protein, and organic substance metabolic processes were significantly induced in Las compared to midrib tissues. In addition, Las in seed coat samples undergoes drastic reprogramming at the RNA and protein level with significant upregulation of genes involved in transcription, translation, protein turnover, and metabolism compared to midrib samples (Supplemental Dataset 2). For example, genes encoding the CarD family transcription factor (*CLIBASIA_01510*) and transcriptional regulator (*CLIBASIA_05645*) were significantly upregulated while *nusA*, encoding a transcription terminator, was significantly downregulated. These represent potential key players in the transcriptional reprogramming only occurred in Las colonizing the seed coat, but not in midrib. Several genes involved in translational machinery were also upregulated in seed coat samples, including those encoding a tRNA synthetase (tilS), t-RNA ligases (*aspS*, *CLIBASIA_03465*, *CLIBASIA_01005*), a translation initiation factor (*CLIBASIA_1775*), and a peptide chain release factor (*CLIBASIA_03855*). Additionally, genes involved in the glutamine metabolism pathway and the TCA cycle converting succinate to oxaloacetate including *fumC*, *CLIBASIA_02745*, *CLIBASIA_4725*, and *mdh* were also upregulated in Las colonizing the seed coats compared to midribs. These findings highlight tissue type-dependent metabolic states of Las in planta. Despite these differences, we found that genes involved in motility such as the flagellar machinery are significantly downregulated in Las during in planta colonization of both tissue types compared to ACP samples (De Francesco et al. 2022).

### The dynamic transcriptional landscape of Las reveals distinct co-regulated genes in planta

To define overall changes in Las gene expression during colonization of the two citrus tissues and identify patterns of co-expressed genes, we analyzed the Las transcriptome using hierarchical clustering. More specifically, hierarchical clustering of the Las genes was performed based on their expression in midrib or seed coat tissues represented as log_2_FC compared to ACP samples. These genes are grouped into seven clusters, each exhibiting a different expression pattern. Clusters I to IV contain genes upregulated in midrib and/or seed coat tissues compared to in ACP whereas clusters V to VII contain downregulated genes (Figure 2A-B). The genes in each cluster were subjected to GO analysis and significantly enriched GO terms were identified (Figure 2C).

**Figure 2.**
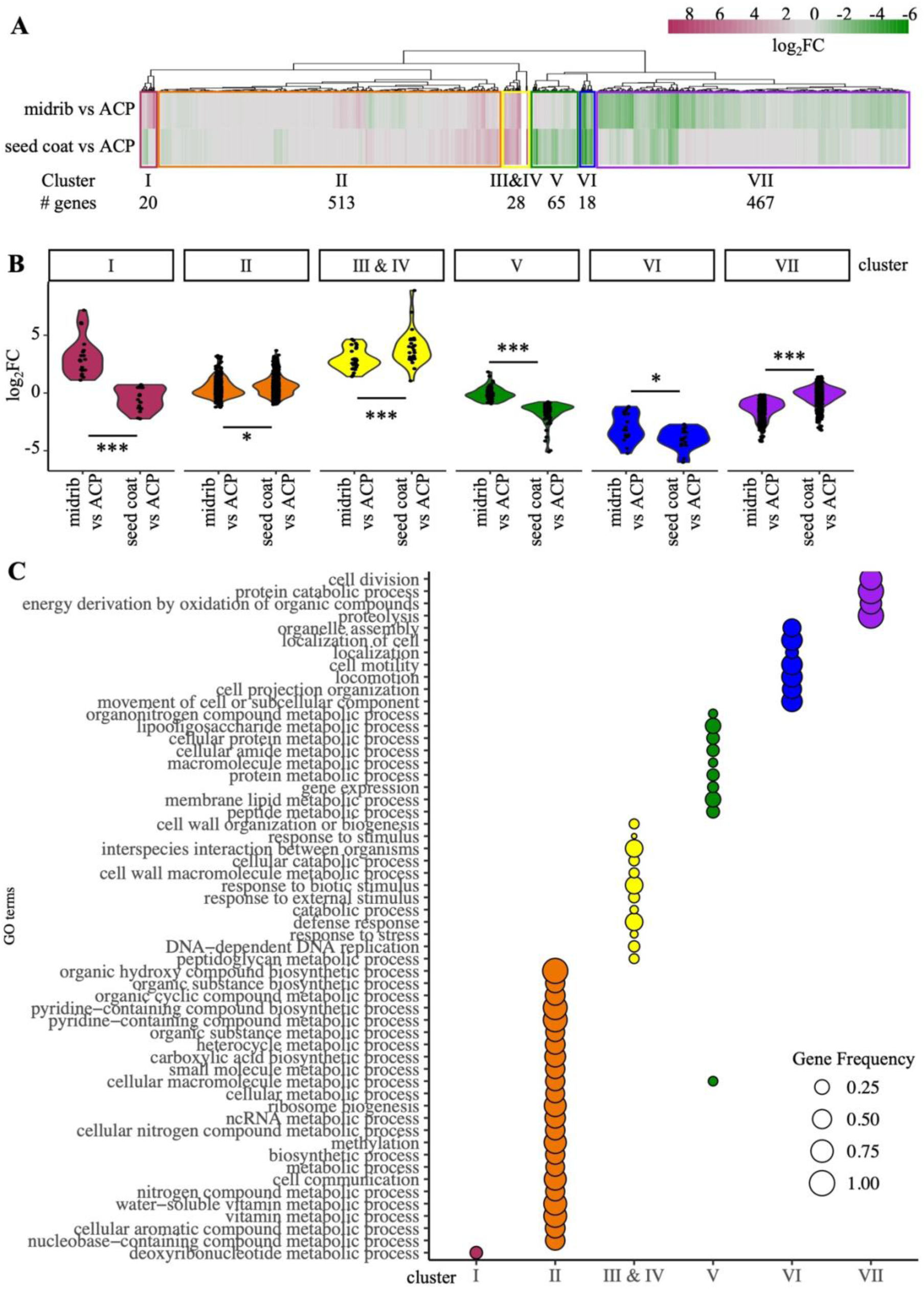
Las exhibits distinct gene expression patterns in different citrus tissue types. **A.** Hierarchical clustering of fold changes in Las gene expression in midrib or seed coat comparing to ACP respectively. Genes were divided into seven clusters based on similar expression patterns where the total number of genes in each cluster is reported. The pheatmap package in R was used for hierarchical clustering and visualization. **B.** Las gene expression displayed as log_2_ fold change (log_2_FC) between midrib and seed coat vs ACP for each gene cluster. Asterisks indicate significant log_2_FC values in midrib or seed coat compared to ACP respectively (2-way ANOVA, * p < 0.05, ** p < 0.01, *** p < 0.001). **C.** Significantly enriched GO terms in each gene cluster. The gene frequency, represented as the number of significantly enriched genes/total number of genes for a given GO term gene group, is indicated by the size of dotplot. For a full GO list, see Supplemental Dataset S3d- j.

Cluster I genes had significantly higher expression in midrib than seed coat and were enriched in genes involved in deoxyribonucleotide metabolism. Cluster II is the largest gene cluster, containing 513 genes, which have similar expression levels in both citrus tissues compared to ACP, but exhibiting significantly higher expression in seed coat than in midrib.

Genes related to various metabolic processes were enriched in this cluster. More specifically, these genes are involved in the biosynthesis of ribosomes, DNA, tRNA, and fatty acids, transcription and translation machinery, and vitamin metabolism. Genes in Cluster III&IV were upregulated in both citrus tissue types vs ACP, with significantly higher expression in seed coat. These genes were found to regulate responses to stress and various stimuli, and defense response. One example of such genes is the phage-related lysozymes, which could play a role in restructuring the bacterial-cell wall during in planta colonization.

Genes grouped in clusters V-VII were repressed in both citrus tissues compared to ACP. Cluster V genes have significantly lower expression in seed coat tissues than in midrib. Genes in this cluster were enriched for lipid and protein metabolism pathways. In particular, these genes encode ribosomal proteins and enzymes involved in lipid polysaccharide biogenesis. Cluster VII genes have significantly higher expression in seed coat than in midrib. These genes are mainly involved in cellular respiration and cell division. Cluster VI has genes that were most downregulated in both citrus tissues compared to ACP. Similarly to our findings in the DEG analysis, these genes were significantly enriched for pathways involved in cell motility and locomotion.

### Transcription factors potentially regulate transcriptome reprogramming

Changes in gene expression is often mediated by binding of specific transcription regulators to specific promoter regions. The Las genome is predicted to encode 19 transcription regulators of which six are found to be essential for viability based on studies of their homologs in the non-pathogenic *Liberibacter crescens* (Lai et al. 2016) and *Sinorhizobium meliloti* (Barnett et al. 2019). Currently, our understanding in how gene expression changes are regulated in Las is very limited. Using the transcriptomic data, we searched the promoter regions (1000 bp upstream of the transcriptional start site) of the genes in each of the seven clusters (Figure 2) for known transcription factor (TF)-binding motifs. Our analysis revealed that eight motifs are significantly enriched in the promoters of the cluster II genes and five in the cluster VII genes (Figure 3A, Supplemental Dataset 4). Only one motif, CodY, was found to be enriched in both gene sets. The unique motifs enriched from each gene set is consistent with the opposite expression patterns of these two clusters. These results suggest that specific TFs associate with the promoter of genes that showed differential expression in different hosts.

**Figure 3.**
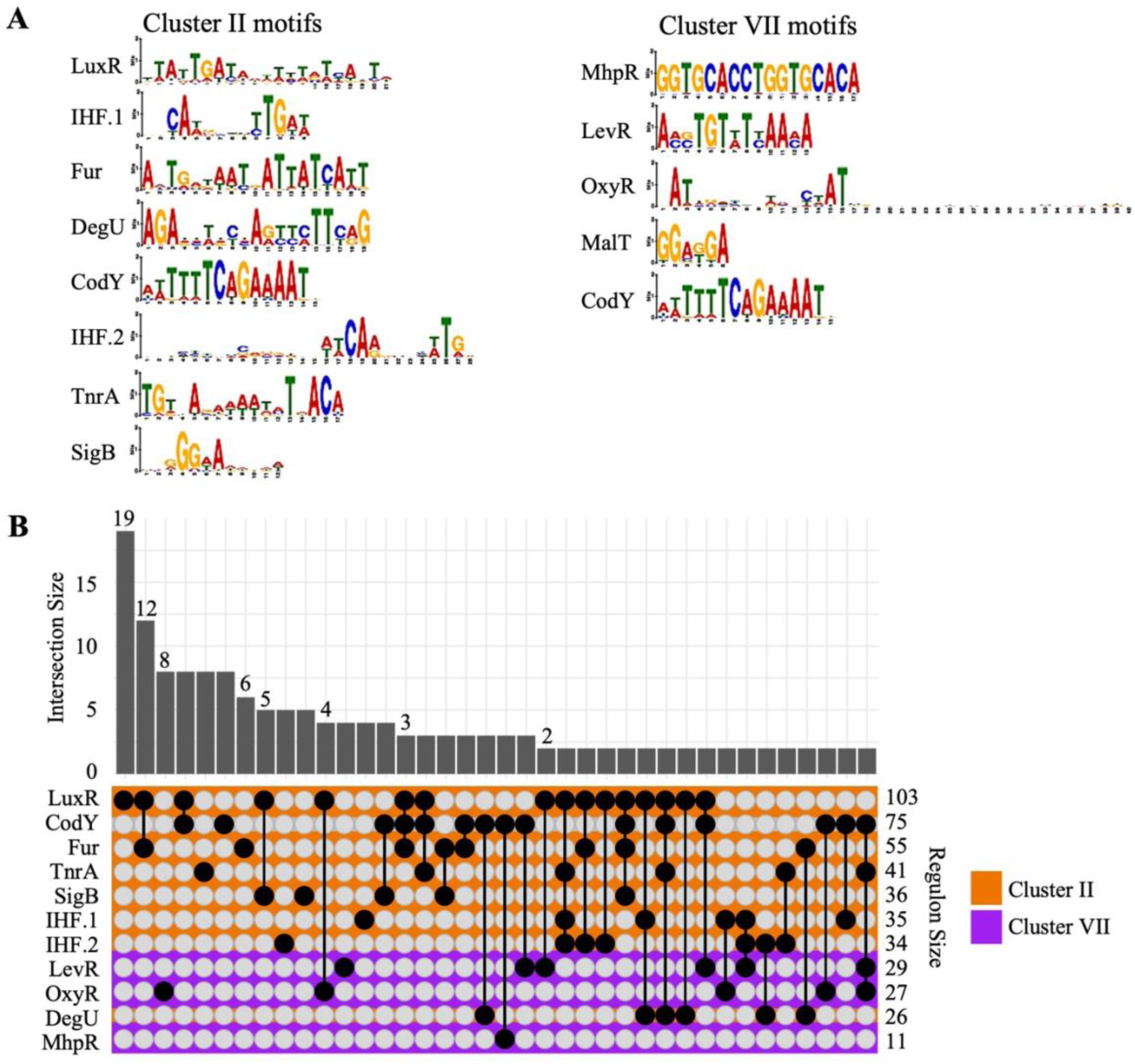
Transcription Factor-binding (TF) motifs enriched in the promoters of Las gene exhibiting differential expression in citrus vs ACP hosts. **A.** Logos of significantly enriched TF-binding motifs in the promoters of genes exhibiting higher expression (Cluster II) or lower expression (Cluster VII) in planta. The gene clusters are presented in Figure 2. Enrichment was determined using 1000 bp region upstream of start codon. Significance was evaluated based on a q-value < 0.05. For a complete list of all the enriched motifs, see Supplemental Dataset S4. **B.** Comparison of downstream Las genes of promoters containing enriched motif occurrences or co-occurrences. Single genes with motif occurrences are not shown. Motifs found to be enriched in Cluster II and VII genes are indicated by orange and purple respectively. Bar plot indicates the number of genes from each interaction. For a full list of motif regulons, see Supplemental Dataset S5a.

We next examined the genes that exhibit higher expression levels in either citrus or ACP to determine which gene promoter contains the above identified motif(s). To understand how specific the enrichment of these motifs is in the genes differentially expressed in citrus vs ACP, we searched each motif using their consensus sequence for occurrences across predicted promoter regions in the entire Las genome and extracted the downstream gene-coding sequences (Supplemental Dataset 5). GO term analyses were subsequently performed to determine the cellular processes that may be regulated by the corresponding TF-binding motifs (Supplemental Figure 1, Supplemental Dataset 3).

Downstream genes of promoters containing the enriched motifs were grouped based on the occurrence of one motif or co-occurrence of multiple motifs (Figure 3B). In general, most of the genes contain more than one motif in their promoter, indicating multiple layers of regulation by different TFs. Fifty-nine genes were found to have unique combinations of motifs in their promoter that was not shared by any other genes in the genome. The largest group of 19 genes has a single LuxR motif, although this motif is also present in an additional 53 genes in combination with other motifs. In particular, 20 of the 53 genes contain the CodY motif in their promoters and an additional 19 contain the Fur motif. Binding motifs of LuxR, CodY and Fur are all enriched in Cluster II gene promoters. Therefore, TFs binding to these motifs potentially work together to reprogram Las transcription during its transition from colonizing the insect to citrus.

We next explored potential TFs in Las that could be involved in regulating genes containing the enriched motifs. Two LuxR-type TFs, CLIBASIA_02900 (visR) and CLIBASIA_02905 (visN), are encoded in the Las genome and have been shown to regulate motility (Andrade and Wang 2019). Indeed, six genes encoding flagellar assembly proteins were predicted to have the LuxR-binding motif in their promoters in our analysis (Supplemental Dataset 5). The 19 genes containing a single LuxR motifs in their promoters encode three hypothetical proteins (CLIBASIA_00420, CLIBASIA_00430, CLIBASIA_00970), two ribosomal proteins, two ligases (CLIBASIA_00850, GlnA), components of DNA replication machinery (CLIBASIA_01695, DnaA), and membrane proteins (CLIBASIA_00995, CLIBASIA_01500, Lnt). Therefore, VisN and VisR may also regulate these cellular processes beyond motility.

The Ferric uptake regulator (Fur) binds to the Fur motifs in the promoters of iron-responsive genes and acts as a repressor in *E. coli* (Escolar et al. 1998). However, Las does not encode a Fur-like TF. Therefore, we looked for other TFs that may potentially bind the Fur motif. In Rhizobia, the IscR-type TF RirA (rhizobial iron regulator) is an iron-responsive repressor that binds to a DNA sequence that shares similarity with the Fur motif (Chao et al. 2005, Díaz-Mireles et al. 2005, Ngok-Ngam et al. 2009, Yeoman et al. 2004). Therefore, Las RirA may bind the Fur motifs. This is further supported by the presence of the Fur motif in the promoter of *rirA* itself, consistent with the role of IscR as a repressor of its own expression in *E. coli* (Schwartz et al. 2001). The Fur motif is present in genes encoding metal transport proteins (*CLIBASIA_02135*, *CLIBASIA_03020*) and responsible for the assembly of Fe-S clusters (*sufE*, *sufA*) (Supplemental Dataset 5). These genes were downregulated in planta (Supplemental Dataset 2). Therefore, RirA may repress iron usage during colonization of citrus tissues and this repression is released in ACP.

The CodY TF is a repressor in Gram-positive bacteria that regulates nutritional stress response and virulence (McDowell et al. 2012, Sonenshein 2005). We found similarly enriched GO terms from genes containing the CodY motif in Las including the Sec secretion machinery, nitrogen compound transport, and pyruvate/carbohydrate metabolism (Supplemental Figure 1, Supplemental Dataset 3). However, the Las genome lacks a CodY family TF. The regulator responsible for binding to this motif remains unknown.

A much smaller set of genes have motifs that were significantly enriched in the cluster VII gene promoters. The largest group has eight genes that have a single OxyR-binding motif (Figure 3B). Most genes in the OxyR regulon do not have other cluster VII-enriched motifs in their promoters, indicating that these regulators likely work independently during ACP colonization. Interestingly, many genes containing the OxyR motif also have other motifs in their promoters including LuxR, IHF.1 and CodY (Figure 3B), which were enriched from clusters with opposing expression patterns. Indeed, 50 gene promoters were found to have at least one cluster II-enriched motif and at least one cluster VII-enriched motif, indicating a network of TFs with complex regulation in Las.

The TF OxyR belongs to the LysR family and regulates genes for oxidative stress defense and DNA repair (Ochsner et al. 2000). LysR family TFs are also important regulators during rhizobial interaction with plant hosts (Schlaman et al. 1992, von Lintig et al. 1994, Habeeb et al. 1991). Las encodes one LysR-type TF, CLIBASIA_00835, which was previously shown to activate a peroxiredoxin gene when expressed in *S. meliloti* (Barnett et al. 2019). We found that genes encoding enzymes related to DNA repair, including DNA topoisomerase (*topA*), DNA exonucleases (*sbcB, CLIBASIA_01760*) and a double-strand break repair protein (*addB*) also have the OxyR motif in their promoters (Supplemental Dataset 5). Similarly to *CLIBASIA_00835*, *topA*, *sbcB* and *addB* were also upregulated in planta, indicating that CLIBASIA_00835 may promote the DNA repair machinery during colonization of citrus (Supplemental Dataset 2). Additionally, genes encoding oxidoreductases, including superoxide dismutase (*CLIBASIA_01910*) and ferredoxin reductase (*CLIBASIA_01240*), also have the OxyR motif in their promoters but were found to be downregulated in planta relative to ACP (Supplemental Dataset 2 & 5). Therefore, the expression of these genes may be mainly regulated by other TFs given that OxyR is known to activate these genes in other bacteria.

### LuxR motif is enriched in promoters of genes encoding virulence factors

As virulence is a tightly regulated process in bacterial pathogens, we sought to determine which of the enriched motifs may regulate virulence gene expression. For this purpose, we searched for occurrences of the significantly enriched motifs from cluster II and cluster VII in 30 known or predicted virulence factor gene promoters using their consensus sequence (Figure 4A, Supplemental Dataset 5). These virulence factors include 26 core SDEs, two phage-encoded peroxiredoxins (Bcp and Prx5), a salicylic acid hydroxylase (SahA), and a putative secreted peroxidase (CLIBASIA_05590). The LuxR-binding motif was found to be the most significantly prevalent in these promoters with 18 sites predicted from the upstream intergenic region of ten SDE gene or gene operons (Figure 4B). These include some of the most highly expressed SDE-encoding genes in planta, such as *CLIBASIA_03230*, *CLIBASIA_03160*, *CLIBASIA_04560*, and *CLIBASIA_04580*. Therefore, LuxR-type TF(s) may regulate virulence gene expression during in planta colonization. In addition, two SDEs, *CLIBASIA_03160* and *prx5*, were found to have significant matches to the Fur-binding motif in their promoters, while *CLIBASIA_04530* had a significant match to the TnrA-binding motif (Supplemental Dataset 5). The SDE-encoding genes *CLIBASIA_03160* and *CLIBASIA_04530* had multiple cluster II motifs in their promoters, suggesting that more than one TF may be involved in their regulation.

**Figure 4.**
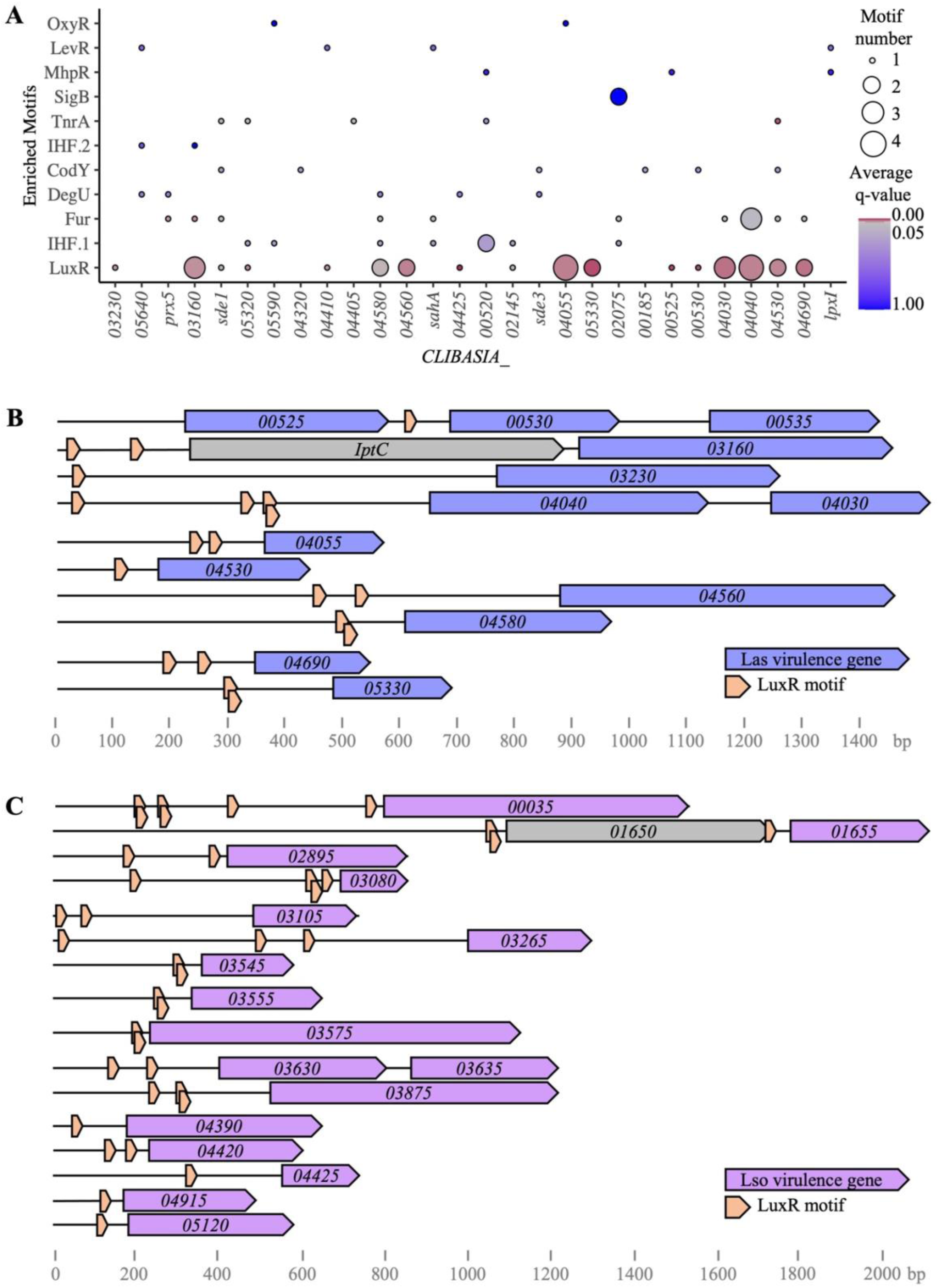
LuxR-binding motif is enriched in the promoters of predicted *Candidatus* Liberibacter spp. virulence genes. **A.** Occurrence of enriched motifs in promoters of Las virulence genes. Consensus sequence of each motif was scanned in the 1000 bp upstream region of all predicted virulence genes in Las genome. The average q-value for motif hits is indicated by the dotplot color. The number of motifs occurrences is indicated by the dotplot size. For full list of individual motif occurrences, see Supplemental Dataset S5b. **B.** Genome regions containing significant (q-value < 0.05) intergenic LuxR motif occurrences upstream of Las virulence genes/operons. Gene names are based on CLIBASIA_ annotation from the Las psy62 genome. **C.** Genome regions containing significant (q-value < 0.05) intergenic LuxR motif occurrences upstream of predicted effector genes/operons in the potato zebra chip pathogen *Ca.* Liberibacter solanacearum (Lso). Gene names are based on CKC_ annotation from the Lso ZC-1 genome. Open reading frames and motifs are color-coded to indicate function. For full list of individual motif occurrences in Las and Lso, see Supplemental Dataset S5c.

To determine whether a similar transcription regulation mechanism is also used in other *Ca.* Liberibacter pathogens, we examined the potato zebra chip pathogen *Ca.* Liberibacter solanacearum (Lso). Lso and Las are closely related bacteria despite colonizing different plant and psyllid hosts. A pairwise comparison between Lso strain ZC-1 and Las strain psy62 reveal high sequence identity across the entire genomes. This conservation extends to pathogenesis-related genes including those that encode lipid polysaccharide synthases, the Sec machinery, and the type I secretion system (Thapa et al. 2020). Lso is also predicted to secrete effectors through the Sec secretion system, called Hypothetical protein effectors (HPEs) (Reyes Caldas et al. 2022). However, their HPE effector repertoire is largely divergent from that of Las.

We searched the LuxR-binding motif in the promoter regions of the 38 predicted HPE genes in the Lso isolate ZC-1 genome and found 39 significant intergenic occurrences upstream of 17 HPE genes (Figure 4C). Ten HPEs containing the LuxR motif in their promoters have detectable expression during Lso infection of tomatoes (Reyes Caldas et al. 2022). It is important to note that none of the 17 HPEs are homologs of the Las SDEs predicted to contain the LuxR motif in their promoters. Nonetheless, the two LuxR-type TFs, VisN and VisR in Las, are conserved in Lso, corresponding to CKC_00545 and CKC_00540 in the ZC-1 genome, respectively. These results strongly indicate a conserve regulatory mechanism for effector expression mediated by the LuxR-type TFs in plant pathogenic Liberibacter.

### Tissue-specific expression of virulence-related genes

Las colonization is not uniform across different citrus tissues, with a much higher bacterial titer in seed coat. Our PCA analysis indicates significant transcriptomic differences between Las colonizing midrib and seed coat tissues, indicative of specific gene expression patterns, and thus host manipulation, that may facilitate the adaptation to distinct host environments. We examined 30 virulence genes including 26 SDEs for their relative expression in seed coat vs leaf midrib (Figure 5A). With the increased coverage of the Las genome using the seed coat tissue, we were able to detect eight virulence factors that were previously undetectable in midrib samples, potentially due to their overall low expression levels. Six of the virulence factors were considered to have “low midrib expression” with their transcripts detectable in only one midrib sample but consistently from seed coat samples. These genes include putative SDEs, including two with known virulence function, *sde3* (Shi et al. 2023) and *sde15* (Pang et al. 2020). The remaining genes were considered to have “high midrib expression” with their transcripts detectable in most midrib and all seedcoat samples. Sixteen SDE-encoding genes were highly expressed in seed coat and/or midrib. In particular, genes encoding the effectors SDE1 and CLIBASIA_03230, which have known virulence functions by targeting the defense-related proteins, papain-like proteases and calcium-dependent kinases, respectively (Clark et al. 2018; Clark et al. 2020, Zhang et al. 2024) were among the most highly expressed genes in planta (Figure 5A). Non-canonical secreted proteins like Prx5, Bcp, SahA, and CLIBASIA_05590 (a putative peroxidase) were also highly expressed in planta. These proteins contribute to virulence by suppressing plant defense responses such as the ROS burst, callose accumulation, and salicylic acid production (Jain et al. 2015, Jain et al. 2018, Li et al. 2017).

**Figure 5.**
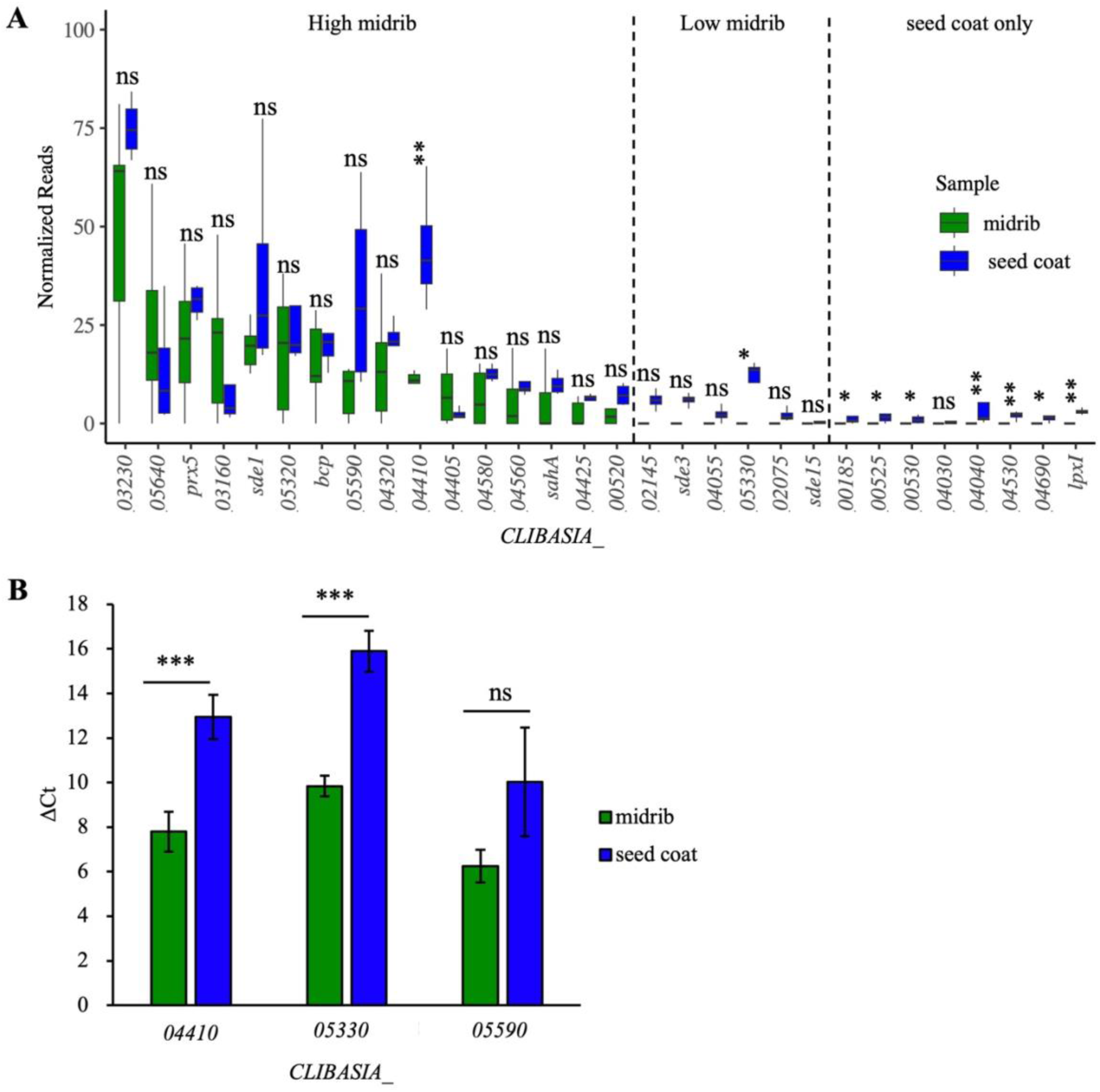
Dynamic expression of Las genes encoding virulence factors in different citrus tissue types. **A.** Different expression of predicted virulence genes in midrib (green) and seed coat (blue) tissue represented by normalized read counts. Genes are displayed in order based on expression levels in midrib and classified as high (>0 reads for at least 2 out of 6 replicates), low (>0 reads for only one replicate), and seed coat only (0 reads for all replicates). Asterisks indicate normalized read counts significantly different in midrib vs seed coat (Wilcox test ns = not significant, * p < 0.05, ** p < 0.01). **B.** qRT-PCR showing expression levels of three Las virulence genes in midribs (green) and seed coat (blue) tissues (n = 3). Las *16S* rRNA was used as the internal control for normalization Using the ΔCt method. Asterisks indicate significant differences (two-tailed *t* test at ns = not significant, *** p < 0.001).

Of the 22 virulence factors with expression detectable in both tissue types, only two exhibited differential expression. These virulence factor genes encode predicted SDEs, CLIBASIA_04410 and CLIBASIA_05330, both of which had significantly higher expression in the seed coat than in midrib (Figure 5A). We confirmed the differential expression pattern of *CLIBASIA_04410* and *CLIBASIA_05330* using RT-qPCR (Figure 5B). In contrast, *CLIBASIA_05590*, encoding a putative peroxidase, showed unchanged expression in both citrus tissues. These results indicate differential expression of these SDEs may result in specific host manipulation in different citrus tissue types.

### Las virulence factors suppress callose deposition

Compared to leaf midrib, Las infected seed coat tissue exhibits reduced ROS accumulation, callose deposition, and/or P-protein production (Bernardini et al. 2022). Given these differences in plant immune responses between midrib and seed coat during Las colonization, we investigated a potential contribution of the virulence factors that exhibited higher expression in seed coat to the enhanced suppression of ROS and callose deposition. For this purpose, CLIBASIA_04410 and CLIBASIA_05330 were examined for suppression of immune-activated ROS burst and callose deposition after transient expression in *Nicotiana benthamiana*. CLIBASIA_05590 was also included as it shares homology with a phage-encoded peroxidase in another Las isolate, which was previously shown to enhance resistance to ROS in *L. crescens* (Jain et al. 2015).

Genes encoding CLIBASIA_04410, CLIBASIA_05330 and CLIBASIA_05590 without the predicted signal peptide were cloned under the control of the CMV 35S promoter and with a C-terminal tag of enhanced GFP (eGFP). These genes were expressed in *N. benthamiana* leaves via agroinfiltration. eGFP and HopF2, a *Pseudomonas syringae* effector with known plant immune suppression activity (Hurley et al. 2014), were used as the negative and positive control, respectively. Infiltrated *N. benthamiana* leaves were treated with the bacterial epitope flg22 or inoculated with the disarmed bacterial pathogen, *P. syringae* pv. tomato DC3000 Δ*hrcC,* to activate immunity and subsequently tested for early (ROS burst) and late (callose deposition) immune outputs. 10 mM MgCl_2_ was used as a mock treatment.

While flg22-triggered ROS burst was not affected by the Las virulence factors (Supplemental Figure 2), a significant reduction in callose deposition, measured by fluorescent areas as the percentage of the total leaf area, was observed in *N. benthamiana* leaves expressing HopF2, CLIBASIA_05590 or CLIBASIA_04410 (Figure 6A-B). Protein accumulation of the eGFP-tagged virulence factors and the eGFP negative control was confirmed via western blotting (Figure 6C). These results suggest that the putative peroxidase, CLIBASIA_05590, and the SDE, CLIBASIA_04410, are able to suppress callose deposition in a ROS-independent manner. Their high expression levels in seed coat may contribute to the enhanced suppression of plant immunity, especially callose deposition, in this tissue type.

**Figure 6.**
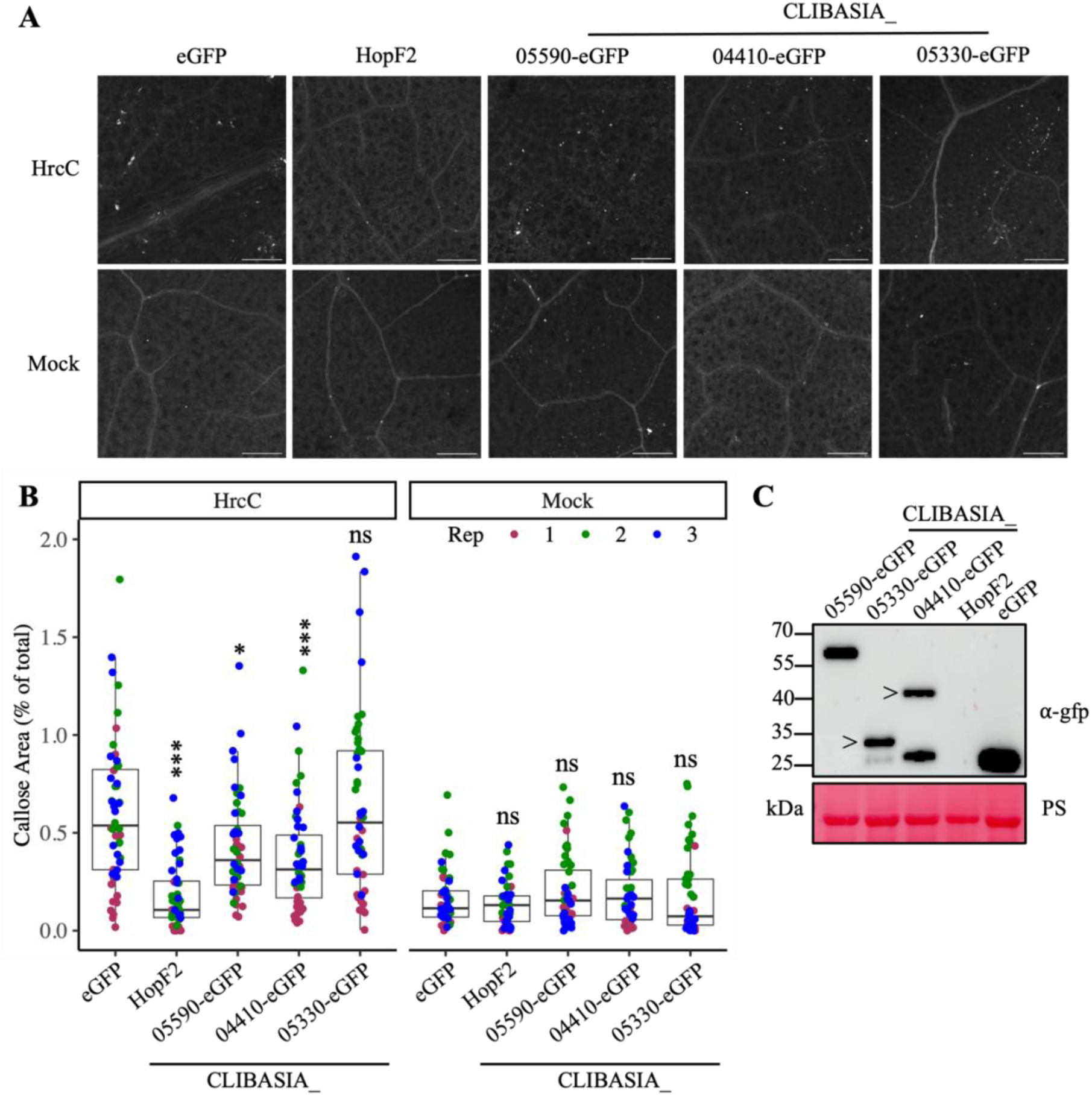
Two Las virulence proteins inhibited bacterial infection-triggered callose deposition when expressed in *Nicotiana benthamiana*. **A-B.** Callose deposition elicited by the bacterium *Pseudomonas syringae* pv. tomato DC3000 Δ*hrcC* (HrcC) was inhibited by CLIBASIA_05590 and CLIBASIA_04410 in *N. benthamiana.* Las genes were transiently expressed in *N. benthamiana* leaves through Agroinfiltration. 48 hours post Agroinfiltration, the leaves were inoculated with HrcC and callose deposition was evaluated at 24 hours post inoculation using fluorescence microscopy after aniline blue staining. eGFP and a *P. syringae* effector HopF2 with known immune suppression activity were used as negative and positive control, respectively. 10 mM MgCl_2_ was used as the mock treatment. Scale bars in the micrographs indicate 250 μm. Quantification of callose deposits was determined by % fluorescent areas in total leaf area using ImageJ. Four images were taken from four plants each and three independent experiments (highlighted by different colored dots). Statistical differences were evaluated based on comparisons to eGFP (2-way ANOVA, ns = not significant, * p < 0.05, *** p < 0.001). **C.** Expression of Las virulence genes in *N. benthamiana* were confirmed by immunoblotting using an anti-GFP antibody. Ponceau Staining (PS) of the membrane was used as a loading control. “>” indicates the correct sized product for samples with multiple bands. HopF2 does not have a tag.

## DISCUSSION

In this study, we provide a more complete Las in planta transcriptome by including samples from citrus sink tissues, namely, the seed coat vasculature. Given the low abundance of bacterial RNAs in total RNA extracts from infected plant tissues, many methods have been developed to either enrich for bacterial cells prior to RNA extraction or enrich bacterial RNAs in total RNA extracts (Lovelace et al. 2018, De Francesco et al. 2022, Nobori et al. 2018). The high Las titer in citrus seed coat vasculature tissues allowed us to obtain high coverage of the Las genome and detect almost all (up to 99%) of the Las genes from infected samples using only total RNA extraction methods. This is a substantial improvement compared to the published Las transcriptomes using midrib samples. For example, we were able to detect the expression of all known and predicted virulence factors, eight of which were undetectable using the midrib tissues. Thus, our study significantly advanced a basic understanding of the Las transcriptome during infection.

Citrus responses to Las infection are variable across different tissue types. The canonical view of citrus responses was mostly gained from analysis of leaf midrib tissues, in which Las infection induces accumulation of ROS, callose, and phloem protein, resulting in phloem plugging and collapse (Etxeberria et al. 2009, Achor et al. 2010, Ma et al. 2022). Interestingly, this is in direct contrast with observations made from infected seed coat, which showed increased phloem pore size and downregulation in the expression of callose synthase (*CsCalS*) and phloem protein 2 (*CsPP2*) genes, which also accumulated a higher bacteria titer (Bernardini et al. 2022). Given these differences between these two tissue types, we pursued to understand Las biology and pathogenicity in the context of these tissue types using transcriptomics. Our RNA-Seq analysis show that Las transcriptomes from the seed coat exhibit unique and distinct changes compared to midrib.

This tissue-specificity of Las gene expression indicates adaptive behavior of Las to colonization of seed coat tissues. Several primary metabolic pathways were significantly upregulated in Las during colonization of seed coat including, the TCA cycle, inositol metabolism, purine/pyrimidine metabolism, and translational machinery such as ribosomal proteins and transfer RNAs. The difference in Las metabolic activity between these tissues could be explained by variations in nutritional availability in phloem tissues during HLB infection. Bacterial manipulation of host metabolite transport has been widely documented for apoplastic (Gentzel et al. 2022) and xylem-colonizing pathogenic bacteria (Chen et al. 2010, Xian et al. 2020); however, this has not been observed for phloem-colonizing bacteria. Differences in phloem metabolite composition including sugars, amino acids, and fatty acids were observed between source and sink tissues during HLB infection (Chen et al. 2022). These changes may reflect metabolic activity differences of Las in seed coat vs midrib. Previously we found metabolic activity was not upregulated during colonization of midribs compared to the psyllid vector (De Francesco et al. 2022). The addition of seed coat transcriptome data demonstrates that Las was even more metabolically specialized when colonizing the seed coat tissue.

Las is found to accumulate to a high level in the vasculature tissue in the seed coat. While we can’t exclude the possibility that Las passively accumulates in seed coat due to the hydrostatic pressure from source to sink tissues, our data supports that these bacteria are actively multiplying in the seed coat vasculature. This is demonstrated by the upregulation of Las genes related to cell wall biogenesis and DNA replication in seed coat tissues. Our data is consistent with transmission electron microscopy analysis showing Las cell division in infected seed coat tissues (Achor et al. 2020).

An improved overall understanding of the Las transcriptome in planta allowed us to further investigate the putative transcriptional regulatory elements Las utilizes during citrus colonization. We analyzed the promoter regions of gene clusters with similar expression profiles for the identification of putative cis-regulatory motifs. Several TF-binding motifs were significantly enriched in both upregulated and downregulated genes in planta and correspond to the regulation of both unique and overlapping cellular processes. Given its reduced genome, Las TFs likely have expanded regulons to facilitate transcriptional reprogramming during ACP and citrus colonization. Additionally, these regulons contain motifs enriched from both upregulated and downregulated genes suggesting that Las TFs could serve as activators or repressors for these regulons depending on the host they colonize.

The Las genome encodes 19 TFs of which a portion of them have been characterized in *A. tumefaciens* (Andrade and Wang 2019), *S. meliloti* (Barnett et al. 2019) and *L. crescens* (Lai et al. 2016). Based on the known function of the TF-binding motifs, we identified putative corresponding Las TFs. The LuxR-binding motifs gave the most significant matches and had the largest regulon in our genome-wide analysis. Las encodes two LuxR-type transcription factors, VisN and VisR, which are found in an operon and were previously characterized to regulate motility by binding to the promoters of the *flp3* pilus gene and *rem*, a response regulator (Barnett et al. 2019, Andrade and Wang 2019). This is supported by our analysis which also found the LuxR motif upstream of flagellar machinery-encoding genes as well as *flp3* and *rem*. In addition, we identify processes that may be regulated by VisN and VisR such as transcription and metabolism (Supplemental Figure 1).

Another interesting motif enriched in genes upregulated in citrus is the Fur motif, which was found upstream of metal-responsive genes. Iron-limitation is a signal used by some phytopathogenic bacteria to indicate entrance into an iron-scarce host, subsequently activating virulence genes (Pandey 2023). We identified RirA as a candidate TF that likely binds to the Fur motif in Las. RirA is downregulated in planta, suggesting that ACP may be a more metal-poor environment than citrus. Las could use metal-sensing to signal a host change from insect to citrus thus reprogramming gene expression to adapt to its new environment.

The LuxR and Fur motifs were enriched from upregulated Las genes in planta, whereas the OxyR motif was enriched in downregulated genes. More specifically, we identified OxyR motifs upstream of genes involved in stress responses. LysR-type family TFs are known to bind to the OxyR motifs. A LysR-type TF in *Pseudomonas aeruginosa* was reported to bind OxyR motifs and positively regulate oxidative stress defense and DNA repair (Ochsner et al. 2000). Las encodes a LysR-type TF, CLIBASIA_00835, which may play a crucial role in response to plant defense response during infection including ROS production. The remaining enriched motifs and their predicted regulons are not further discussed as they either lack corresponding regulator homologues in Las or they are likely bound by essential regulators for viability. Further investigation is needed to verify our prediction of Las TF binding to these gene regulons.

Virulence is a tightly regulated process in pathogenic bacteria where the coordinated deployment of virulence factors is essential for pathogenicity once the bacteria enter their host (Lovelace et al. 2023). Several regulatory motifs and their corresponding master regulators have been identified for virulence gene regulons in phytopathogenic bacteria but not phloem-colonizing bacteria (Mole et al. 2007). In phytopathogenic bacteria, most virulence genes are induced in the presence of the host and/or quorum sensing molecules (Wang et al. 2015). Similarly, *Ca.* Liberibacter spp. are known to have different expression patterns between plant and psyllid hosts (De Francesco et al. 2022, Reyes Caldas et al. 2022). Our transcriptomic data allowed us to identify LuxR as an enriched motif in the promoters of predicted effectors in both Las and Lso. LuxR motifs are known to be bound by the LuxR-type TFs. VisN and VisR are conserved LuxR-type TFs among Rhizobia that potentially regulate effector gene expression. This suggests that virulence regulation mechanism is conserved among *Ca.* Liberibacter spp. despite a lack of conservation in the effector genes themselves.

As VisN and VisR are downregulated in planta relative to ACP, it is possible that they serve as negative regulators for the effector genes. In other Gram-negative bacteria, LuxR is also a key component of the LuxI-LuxR system. The *luxI* gene encodes N-acyl-homoserine lactone (AHL) synthase, which is the quorum sensing signal that regulates motility, virulence, and survival (Baltenneck et al. 2021). Las *visN and visR* are considered “solo” *luxR* genes because they lack a corresponding *luxI* gene (Hudaiberdiev et al. 2015). Therefore, they may mainly respond to non-quorum sensing signals to regulate transcription.

Further studies are needed to determine if these LuxR-type transcription factors indeed bind to these motifs and regulate Las virulence gene expression in citrus.

The different responses in ROS accumulation and callose deposition in seed coat tissues may be explained by tissue type-specific manipulation by Las. We compared virulence genes expression in seed coast vs midrib and identified two putative SDEs, with significantly higher expression in seed coat. One of these SDEs, CLIBASIA_04410, encoding a hypothetical protein, could suppress bacteria-induced callose deposition after transient expression in *N. benthamiana* in a ROS-independent manner. In addition, another gene *CLIBASIA_05590*, which is also highly expressed in seed coat, also suppressed callose deposition. CLIBASIA_05590 shares homology with the functional peroxidase SC2_gp095 encoded by a bacterial phage (Jain et al. 2015). Although we did not observe suppression of the PTI-triggered ROS burst by either CLIBASIA_05590 or CLIBASIA_04410, these virulence proteins could still affect the prolonged H_2_O_2_ accumulation during HLB infection in citrus. Further investigation is required to determine the mechanism by which these virulence factors suppress plant defense.

In summary, the current study makes a significant contribution to the knowledge of HLB pathogenesis and provide novel insights into the basic biology of Las including adaptation to different host environment and regulatory components that contribute to transcription reprogramming.

## MATERIALS AND METHODS

### Plant materials and bacterial strains

Seed coat tissues were collected from Duncan grapefruit (*Citrus paradisi*). Four biological replicates of HLB-infected Duncan grapefruit were taken from symptomatic trees in a field heavily exposed to HLB in Florida. Healthy samples from trees grown under protective screens were taken as controls. Vasculature was isolated as described in (Bernardini et al. 2022). Las infection was confirmed, and their titer was determined by reverse transcription-quantitative polymerase chain reaction (RT-qPCR) using Las 16S primers (Yan et al. 2013). The Ct values of the Las-infected samples were compared with those from the healthy tissues as controls.

*N. benthamiana* plants were grown in a controlled growth room with 16:8 light:dark conditions at 22°C, 80% humidity. Five-week-old *N. benthamiana* plants were used for agroinfiltration.

Strains and plasmids used in this study are listed in Table S1. *Pseudomonas* strain used in this study is the Type III Secretion mutant *P. syringae* pv. tomato DC3000 Δ*hrcC* (Yuan and He 1996). *Agrobacterium tumefaciens* strain GV3101 was used for *Agrobacterium*-mediated transient expression in *N. benthamiana*.

### RNA extraction

For RNA extraction from the citrus midrib samples for quantitative reverse transcription PCR (RT-qPCR), 1 mL of TLE buffer (200 mM Tris-HCl (pH 8.2), 100 mM LiCl, 50 mM EDTA, 10% SDS, 2% β-mercaptoethanol) was added to 100 mg citrus powder. 60 µL of 500 mM CaCl_2_ was added to the suspension or the bacterial cells isolated from the citrus tissue as described above, mixed thoroughly, and placed on ice for 10 min. The solution was centrifuged at 12,000 x g for 10 min at 4°C, followed by two rounds of extraction using 500 µL of phenol:chloroform:isoamyl alcohol (25:25:1) and centrifugation at 15,000 x g for 10 min at 4°C. The supernatant was mixed with 2 M LiCl and incubated overnight at - 20°C. RNA was precipitated by centrifugation at 15,000 x g for 30 min, washed twice with ice-cold 75% ethanol, and re-suspended in 20 µL of RNAse-free water. The RNA extracts were treated with Dnase I (Thermo Fisher Scientific) to remove genomic DNA.

For RNA extraction from healthy and diseased citrus seed coats for RNASeq and RT-qPCR, three 100 mg of seed coat vasculature samples were ground under freezing conditions using a GenoGrinder and tungsten beads at 1500 rpm for 30 seconds twice. The RNA was extracted from the homogenized tissue using the Qiagen RNeasy Plant Mini Kit according to the handbook instructions which included an on column Dnase I treatment. Samples were eluted with 50 uL of RNase-free water. A fourth diseased seed coat sample had RNA extracted using the same bacterial isolation methods as described in De Francesco et al. (2022).

The RNA concentration and quality of the extracts were determined by agarose gel, nanodrop and an Agilent 2100 Bioanalyzer using the eukaryotic analysis software.

### Quantitative reverse transcription PCR

We examined the effectiveness of the bacterial RNA enrichment from midrib and seed coat samples by comparing the expression levels of Las *16S* rRNA (Yan et al. 2013) and a f-box gene (Citrus Unigene ID CAS-PT-306416) from Poncirus trifoliata (Mafra et al. 2012) using RT-qPCR. Primer sequences are listed in Supplemental Table 1. RT-qPCR procedure for midrib citrus samples were conducted as described in De Francesco et al. (2022). For citrus seed coat samples, cDNA was synthesized from 1 µg of total DNase-treated RNA using the QuantiTect Reverse Transcription Kit (Qiagen) in 20 µL final volume according to manufacturer’s instructions. qPCR reactions were performed in a CFX96 real-time PCR detection system (Bio-Rad) containing 100 ng of cDNA template (2 µL), ChamQ Universal SYBR qPCR Mast Mix (Vazyme), RNase-free water and 10 µM of gene-specific primers in a final volume of 10 µL. Cycling conditions were: 95°C for 2 min, followed by amplification with 39 cycles of 95°C for 15 sec, 54°C for 30 sec, and 72°C for 30 sec. Relative abundance of the Las RNA was calculated using the equation ΔCt = Ct_Las 16S_ - Ct_Citrus Fbox_ where the Ct values were the average of three technical replicates.

RT-qPCR for virulence factor candidates in citrus was performed using the same procedure described above and primers specified in Supplemental Table 1. The QuantiTect SYBR Green RT-PCR kit (Qiagen) was used to synthesize the cDNA and perform qPCR under the following conditions: 95°C for 30 sec and then 40 cycles of 95°C for 10 sec, 56°C for 30 sec, and 72°C for 30 sec including a melt curve. Virulence factor differential expression was determined using the formular ΔCt = (Ct_Las vir_ – Ct_Las 16S_) where the Ct values of three technical replicates were averaged and the relative expression of 3 biological replicates were calculated individually for each tissue type so that virulence factor and reference gene values could be paired appropriately for data collection within runs. For each virulence gene, a t-test for pairwise comparison (p < 0.05) was used to determine whether expression of the virulence factor was significant between citrus tissue types for three biological replicates per sample type.

### RNA-seq

The seed coat RNA libraries were constructed using either TruSeq Stranded Total RNA Library Prep Kit (Illumina) or Meta-transcriptome Library (Novogene). The RNA samples were treated using the Illumina Ribo-Zero Plus rRNA depletion kit as part of the standard protocol. cDNA libraries were then subjected to RNA-seq using either an Illumina HiSeq-4000 system with 100-bp strand-specific single-end reads or IlluminaNovaSeq-6000 system with 150-bp strand-specific paired-end reads. Library preparation and sequencing were performed in the DNA Technologies Core Genome Center at University of California, Davis or Novogene.

### RNA-seq analysis

The quality of the raw sequencing data was checked with FastQC v0.11.9 (Andrews 2010). Subsequent statistical analyses were performed using R and Python. To ensure high-quality sequences for mapping and downstream analyses, low quality reads and adapter had been trimmed by the sequencing facility prior to sequence delivery. RNA-seq reads were aligned with the indexed Las-Psy62 genome assembly (NC_012985.3) and parsed to it using HISAT2 v2.2.1. The number of reads mapped to each gene was counted using the feature counts function of the Rsubread package (Liao et al. 2019) and listed in Supplemental Data Set 1. The seedcoat data used for the analysis has been deposited into the GEO database (accession No. 270557). The midrib and ACP data used for the analysis were previously published GEO database accession No. 180624 (De Francesco et al. 2022).

### Differential Gene Expression analysis

Gene expression values were normalized to counts per million (CPM). Differentially expressed genes (DEGs) were identified from gene counts for each sample using the DESeq2 package v1.30.1 from Bioconductor (Love et al. 2014). DEGs were selected based on log2-transformed and normalized counts that have an adjusted p value (padj) below a false discovery rate (FDR) cutoff of 0.05. Principal components analysis (PCA) of rlog transformed read counts was performed for all treatments and replicates using the plotPCA function in DESeq2. The rlog function transforms the count data to the log2 scale in a way that minimizes differences between samples with low counts. Hierarchical clustering and Euclidean distance clustering were performed on log_2_ fold change values of in planta samples compared to ACP samples using the dist function and pheatmap function of the pheatmap library v1.0.12.

### Motif enrichment analysis

Promoters defined as 1,000 bp upstream of genes in the *Ca.* Liberibacter spp. genomes were extracted using extract-promoter-sequences code downloaded from Github (https://github.com/KTMD-plant/extract-promoter-sequences) using the NCBI gff and fasta files as input for the Las psy62 (NC_012985.3) and Lso ZC-1 (NC_014774.1) genomes. Fasta files of promoters for groups of Las genes based on in planta expression clustering were used as input files for motif enrichment using Simple Enrichment Analysis v.5.4.1 against all prokaryote DNA motif databases (Bailey and Grant 2021). Significantly enriched motifs were identified based on an FDR q value cut-off of 0.05.

### Motif scanning analysis

The consensus sequence of significantly enriched motifs identified from Las gene promoters were used as inputs for motif scanning within the Las genome promoters and Las virulence factor promoters using Find Individual Motif Occurrences v.5.4.1 using the given strand only (Grant et al. 2011). This same method was used to scan for LuxR motifs within the Lso promoters of predicted effectors identified by Reyes Caldas et al. (2022). Significantly occurring motif matches were identified based on an FDR q value cut-off of 0.05. The significant motif occurrences from predicted Las and Lso virulence factor promoters were manually filtered based on whether motifs were intergenic and upstream of the virulence factor gene or gene operon. To determine if candidate motifs were intergenic, their sequence hit was manually mapped to their respective genomes. Gene operon prediction for Las and Lso gene pairs were determined using MicrobesOnline Operon Predictions (Price et al. 2005) and Computational Genomic Group Operon-mapper (Taboada et al. 2018) respectively using NCBI genome inputs. The resulting GFF file output from the motif scan of the Las genome promoters was reformatted to include genomic coordinates and chromosome name based on the NCBI genome. Intergenic motifs from this GFF file were identified using the bedtools v.2.28.0 intersect command using forced strandedness and the NCBI Las genome GFF input file with an overlap bp output (Quinlan and Hall 2010). Motifs with 0 bp overlap were extracted as intergenic and their downstream genes were used as input for gene ontology enrichment analysis.

### Gene Ontology (GO) analysis

Relevant GO terms were recruited manually from the KEGG gene database, where we determined the IDs of the associated proteins in Uniprot DataBase and extracted their complete GO annotations from QuickGO version 2021-02-08. The GO-term analysis was conducted using the biocManager package, ViSEAGO v.1.4.0. The custom GO-terms for Las were annotated using the Custom2GO and its annotation function. Enriched GO-terms were identified from gene sets from each cluster, DEG comparison, or motif outputs using the entire genome as the background gene set using the “create_topGO data” function. To test for significant enrichment, the classic algorithm and fisher exact test were used with a cutoff p value of 0.05. Redundant enriched GO terms were identified using the REVIGO web server using the small parameter (Supek et al. 2011).

### Reactive Oxygen Species (ROS) assay

ROS production was monitored using a luminol-based assay (Keppler et al. 1989). *A. tumefaciens* GV3101 with various constructs were streaked from glycerol stocks onto LBA plates containing appropriate antibiotics. Two-day old bacterial lawns were resuspended in sterile distilled water and washed twice by spinning at 4000 rpm for 10 minutes. The cells were resuspended in MgCl_2_-2-(N-morpholimo)ethanesulfonic acid (MES) buffer (10 mM MgCl_2_, 10 mM MES, pH 5.6) to a final concentration of OD_600_=1.0 before adding acetosyringone to a final concentration of 150 μM. Bacteria were incubated at room temperature in the dark shaking at 30 rpm for three hours before combining in a 1:1 ratio with the strain carrying the viral RNA silencing suppressor P19, meaning the final OD_600_ of each strain was 0.5. Leaves of four five-week old *N. benthamiana* plants were infiltrated with a blunt syringe. Two days after infiltration, leaf disks (diameter of 4 mm) from the infiltrated area were removed using a circular borer. Leaf disks were pre-incubated overnight in 96-well flat bottom, chimney well, LUMITRAC plates (Grenier Bio-One, Austria) with 100 μL of distilled water. The water was discarded and replaced with 100 μL of 10 mM MOPS-KOH buffer (10 mM MES, 10 mM MgCl_2_, 10 mM CaCl_2_, 5 mM KCl_2_, pH 5.8) containing 0.5 mM luminol (Sigma-Aldrich, Unites States, A8511-5g) and 10 μg/mL horseradish peroxidase (Sigma-Aldrich, P6782). ROS was elicited with 100 nM flg22 (Genscript Biotech Corporation, China) or water for the mock controls.

Immediately after treatment, luminescence was measured every two minutes for an hour in a SpectraMax iD5 plate reader (CA, Unites States). Each experiment was repeated three times independently.

### Callose deposition assay

Callose deposition in 10 mm leaf disks was visualized 72 hours after agro-infiltration of four five-week-old *N. benthamiana* plants following aniline blue staining as previously described with modifications (Nguyen et al. 2010). Callose deposition was elicited 48 hours after agro-infiltration by infiltrating *P. syringae* pv. tomato DC3000 Δ*hrcC* suspended in 10 mM MgCl_2_ buffer to a final concentration of OD_600_=0.2 using 10 mM MgCl_2_ buffer as a mock control. The cleared samples were incubated overnight in aniline blue stain, washed, and viewed under a Leica SP8 fluorescent microscope with 10X dry objective. Fluorescence was observed under ultraviolet light (wavelength = 405 nm). Four images were taken from different fields of view for each sample. The callose area was measured using ImageJ software with Trainable Weka Segmentation plugin as previously described (Mason et al. 2020). Each experiment was repeated three times independently.

### Western blotting

To evaluate Las virulence factor expression, 1/4 cm^2^ of tissue was excised across 4 replicate *N. benthamiana* plants for a total of 1 cm^2^ of tissue for each sample. Leaf discs were immediately frozen in liquid nitrogen and stored at -80 °C. Leaf disks were ground under freezing conditions using 10 mm tungsten beads (Qiagen) in the Genogrinder at 1000 rpm for one minute. To extract proteins, 100 μL of protein extraction buffer (50 mM Tris-HCl (pH 7.5), 150 mM NaCl, 1 mM EDTA, 0.1% Triton X-100, 10mM DTT, 1x cOmplete EDTA-free protease inhibitor cocktail (ThermoFisher)) was added to the ground tissue. Crude plant material was removed by centrifugation at 15,000 x g for 15 minutes and 60 μL of the supernatant was combined with 20 μL mPAGE sampling loading buffer (Merck Life Science) and boiled at 95 °C for 5 minutes. Protein samples were separated by SDS-PAGE and immunoblotting was conducted using anti-GFP at a concentration of 1:1000 (Sigma-Aldrich), followed by secondary anti-mouse-HRP at a concentration of 1:5000 (Biorad). Positive signals were detected via chemiluminescence using Super Signal West Femto chemiluminescent substrates (Pierce) and visualized using the Amersham ImageQuant 800 (cytiva).

## AUTHOR CONTRIBUTIONS

1. W. Ma conceived and designed the experiments. C. Wang and A. Levy provided materials. A. H. Lovelace conducted all the experiments including the bioinformatics and statistical analyses. A. H. Lovelace prepared figures and tables and wrote the manuscript with W. Ma. All authors contributed to the article and approved the submitted version.

## DATA AVAILABILITY

Transcriptome data used for analysis have been deposited in the NCBI GEO database (accession no. 180624 and 270557). All other raw data are available upon request to W. Ma.

## ACKNOWLEDGEMENTS

We thank the members of the Ma lab for their helpful discussions and feedback and USDA-NIFA for supporting this research.

