## Supplemental for "Transcriptomic profiling of ‘*Candidatus* Liberibacter asiaticus’ in different citrus tissues reveals novel insights into Huanglongbing pathogenesis"

SUPPLEMENTAL MATERIALS (5 Datasets, 2 Tables, 3 Figures)

**Dataset S1.** Normalized counts of 1111 genes in *Candidatus* Liberibacter asiaticus (Las) transcriptome for bacterial samples collected from the midribs or seed coats of citrus plant host as well as insect vector host, Asian Citrus Psyllid (ACP).

**Dataset S2.** Differentially expressed genes (DEGs) of Las when colonizing of citrus midribs, citrus seed coats, and ACP hosts.

**Dataset S3.** Significantly enriched Gene Ontology (GO) terms found in Las DEGs using pairwise comparisons between different host or tissue types, gene clusters based on expression in citrus vs ACP hosts, and genes containing Transcription Factor-binding (TF) motifs in their promoters.

**Dataset S4.** TF-binding motifs enriched in the promoters of Las gene clusters based on expression in citrus vs ACP hosts.

**Dataset S5.** Individual TF-binding motif occurrences found in Las gene promoters (1000 bp), Las virulence factor gene promoters, and *Candidatus* Liberibacter solanacearum ZC-1 (Lso) effector gene promoters.

**Table S1.** Bacterial strains and constructs used in this study.

**Table S2.** Primers and geneblocks used in this study.

**Figure S1.** Significantly enriched GO terms of downstream Las genes containing enriched TF-binding motifs in their promoters. Dotplot color of motifs found to be enriched in Cluster II and VII genes are indicated by orange and purple respectively. The gene frequency, represented as the number of significant genes/total number of genes for a given GO term gene group, is indicated by the dotplot size. For a full GO list, see Supplemental Dataset S3k-t.

**Figure S2.** Flg22-elicited reactive oxygen species (ROS) burst is unaffected by Las virulence factors when expressed in *Nicotiana benthamiana*. **A-B.** ROS elicited by flg22 was unaffected by Las virulence factors in *N. benthamiana*. Las genes were transiently expressed in *N. benthamiana* leaves through Agroinfiltration. 48 hours post Agroinfiltration, the leaves were treated with 100 nM flg22 (Flg22, circle) or water (Mock, triangle) and total ROS production was evaluated over 60 minutes using a luminol-chemiluminescence assay. eGFP and a *P. syringae* effector HopF2 with known immune suppression activity were used as negative and positive control, respectively. 10 mM MgCl2 was used as the mock treatment. Shaded regions represent mean ± standard deviation of four disks taken from four plants each. Quantification of the cumulative RLU was determined by measured area under the ROS burst curve. Measurements taken from 4 leaf disks from four plants each and three independent experiments (highlighted by different colored dots). Statistical differences were evaluated based on comparisons to eGFP (2-way ANOVA, ns = not significant, *** $p < 0.001$). **C.** Expression of Las virulence genes in *N. benthamiana* were confirmed by immunoblotting using an anti-GFP antibody. Ponceau Staining (PS) of the membrane was used as a loading control. “>” indicates the correct sized product for samples with multiple bands. HopF2 does not have a tag.

**Supplementary References.**

**Table S1.** Bacterial strains and constructs used in this study.

| Strain/Construct | Purpose | Reference |
|---|---|---|
| *Pseudomonas syringae* pv. tomato DC3000 Δ*hrcC* | PTI elicitor, $Rif^R$ | Yuan and He 1996 |
| *Agrobacterium tumefaciens* GV3101 | Agroinfiltration, $Rif^R$, $Gm^R$ | Holsters et al. 1980 |
| GV3101:P19 | Silencing suppressor, $Kan^R$, $Rif^R$, $Gm^R$ | Baulcome and Molnar 2004 |
| pEG100 | Transient expression of tagless effector, $Chlor^R$, $Kan^R$ | Earley et al. 2006 |
| pEG100:HopF2 | Transient expression of tagless effector, $Kan^R$ | This study |
| FP08018-BsaI | Transient expression of eGFP-tagged virulence factor, $Kan^R$ | Reyes Caldas et al. 2022 |
| FP08024 | Transient expression of eGFP, $Kan^R$ | Reyes Caldas et al. 2022 |
| FP08018:5590 | Transient expression of eGFP-tagged virulence factor, $Kan^R$ | This study |
| FP08018:4410 | Transient expression of eGFP-tagged virulence factor, $Kan^R$ | This study |
| FP08018:5330 | Transient expression of eGFP-tagged virulence factor, $Kan^R$ | This study |

**Table S2.** Primers and geneblocks used in this study.

| Name | Organism | Target/gene | Sequence (5' to 3') | Purpose | Reference |
|---|---|---|---|---|---|
| CitrusFboxqPCR.for | *Poncirus trifoliata* | F-box | TTGGAAACTCTTTCGC CACT | qRT PCR | Mafra et al. 2012 |
| CitrusFboxqPCR.rev | *P. trifoliata* | F-box | CAGCAACAAAATACCC GTCT | qRT PCR | Mafra et al. 2012 |
| Clas16SqPCR.for | *Candidatus* Liberibacter asiaticus (Las) | *CLIBASIA_r05781* | GGATAACGCATGGAAA CGTGTGCT | qRT PCR | Yan et al. 2013 |
| Clas16SqPCR.rev | Las | *CLIBASIA_r05781* | AATCCAACGCAGGCTC ATCTCTCT | qRT PCR | Yan et al. 2013 |
| LasgyrBqRT.for | Las | *CLIBASIA_03525* | TTGAACAAGCTGTAATT TCTGG | qRT PCR | Fleites et al. 2014 |
| LasgyrBqRT.rev | Las | *CLIBASIA_03525* | ATCTGTTTGCCAATTTA GAAGC | qRT PCR | Fleites et al. 2014 |
| CLas05590qRT.for | Las | *CLIBASIA_05590* | GGATCCCAAGAGGCTG ATA | qRT PCR | This study |
| CLas05590qRT.rev | Las | *CLIBASIA_05590* | CCCAGACAAAACCCTT GATC | qRT PCR | This study |
| SDE25qRT.for | Las | *CLIBASIA_04410* | CACTGTCTGCGGAAAA TGAA | qRT PCR | Thapa et al. 2020 |
| SDE25qRT.rev | Las | *CLIBASIA_04410* | GTAGCGGTGTCCGTTG TTTT | qRT PCR | Thapa et al. 2020 |
| SDE39qRT.for | Las | *CLIBASIA_05330* | TATAACGTGCGGTGCA CAAG | qRT PCR | Thapa et al. 2020 |
| SDE39qRT.rev | Las | *CLIBASIA_05330* | AAAGAATCAAGATTTTC TTTTGCTG | qRT PCR | Thapa et al. 2020 |
| attL1hopF.for | *Pseudomonas syringae* pv. tomato DC3000 | *hopF* | | cloning into pEG100 | This study |
| attL2hopF.rev | *P. syringae* pv. tomato DC3000 | *hopF* | | cloning into pEG100 | This study |
| BsaI5590.for | Las | *CLIBASIA_05590* | TGGTCTCAAATGGTCG AATGCGAGCGTTC | cloning into FP08018 | This study |
| BsaI5590.rev | Las | *CLIBASIA_05590* | TGGTCTCACGAACTTT GATTAAGTGTTGCTTG CTTTT | cloning into FP08018 | This study |
| BsaISDE25.for | Las | *CLIBASIA_04410* | tggtctcaaatgATAAATCCT TGTGGAATT | cloning into FP08018 | This study |
| BsaISDE25.rev | Las | *CLIBASIA_04410* | tggtctcacgaactGTAACGA TGCCCATTA | cloning into FP08018 | This study |

| | | | | | |
|---|---|---|---|---|---|
| BsaISDE39.for | Las | *CLIBASIA_05330* | TGGTCTCAAATGATATCTTTCCTTCCACAAATTTTC | cloning into FP08018 | This study |
| BsaISDE39.rev | Las | *CLIBASIA_05330* | TGGTCTCACGAACTATATGTTGCAAAAAAAGAATCAAGAT | cloning into FP08018 | This study |
| At_mSDE25 | Codon optimized for *Arabidopsis thaliana* expression and no signal peptide | *CLIBASIA_04410* | CCGGGTCTCCTCGAGATGATAAATCCTTGTGGAATTGAAGAAGACAATTTGAAGTCTTCACCGCTGCCGCATATCGCACTGGAGTCTTTAAGTGCCGAGAACGAGAAGAAGGAGCTTAGCGAGCATGAAAAGAAGGTTATCGAGAGCCAGGAGAACCCTAAGAAGCAGTTTAGTGAGCATGAAAAGAAGGAAACCGATGACCCTAAGAGTGCTAGGAAGGAGAACATCGTCATGAAGAAAACTTTCTCTCAGAAGTCTAAGAAGTATACCCCTTATTTCGATCATTACATGACAAATGGCCACCTGAACTTGCCCCAGAATAATGGGCATCGTTACGGCGGCGGCGGCTCTGGCCGGAGACCG | cloning into FP08018 | This study |

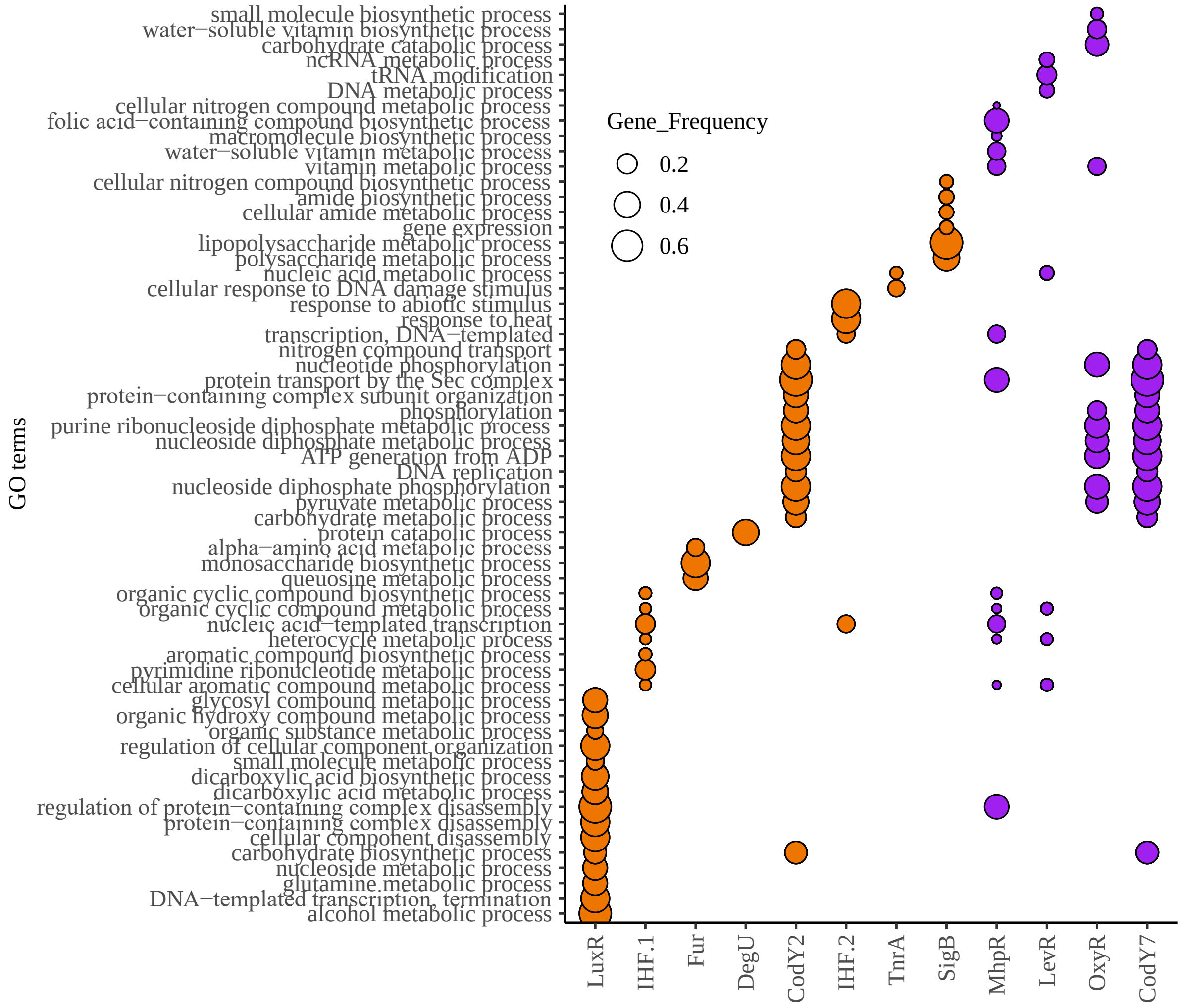


**Figure S1.** Significantly enriched GO terms of downstream Las genes containing enriched TF-binding motifs in their promoters. Dotplot color of motifs found to be enriched in Cluster II and VII genes are indicated by orange and purple respectively. The gene frequency, represented as the number of significant genes/total number of genes for a given GO term gene group, is indicated by the dotplot size. For a full GO list, see Supplemental Dataset S3k-t.

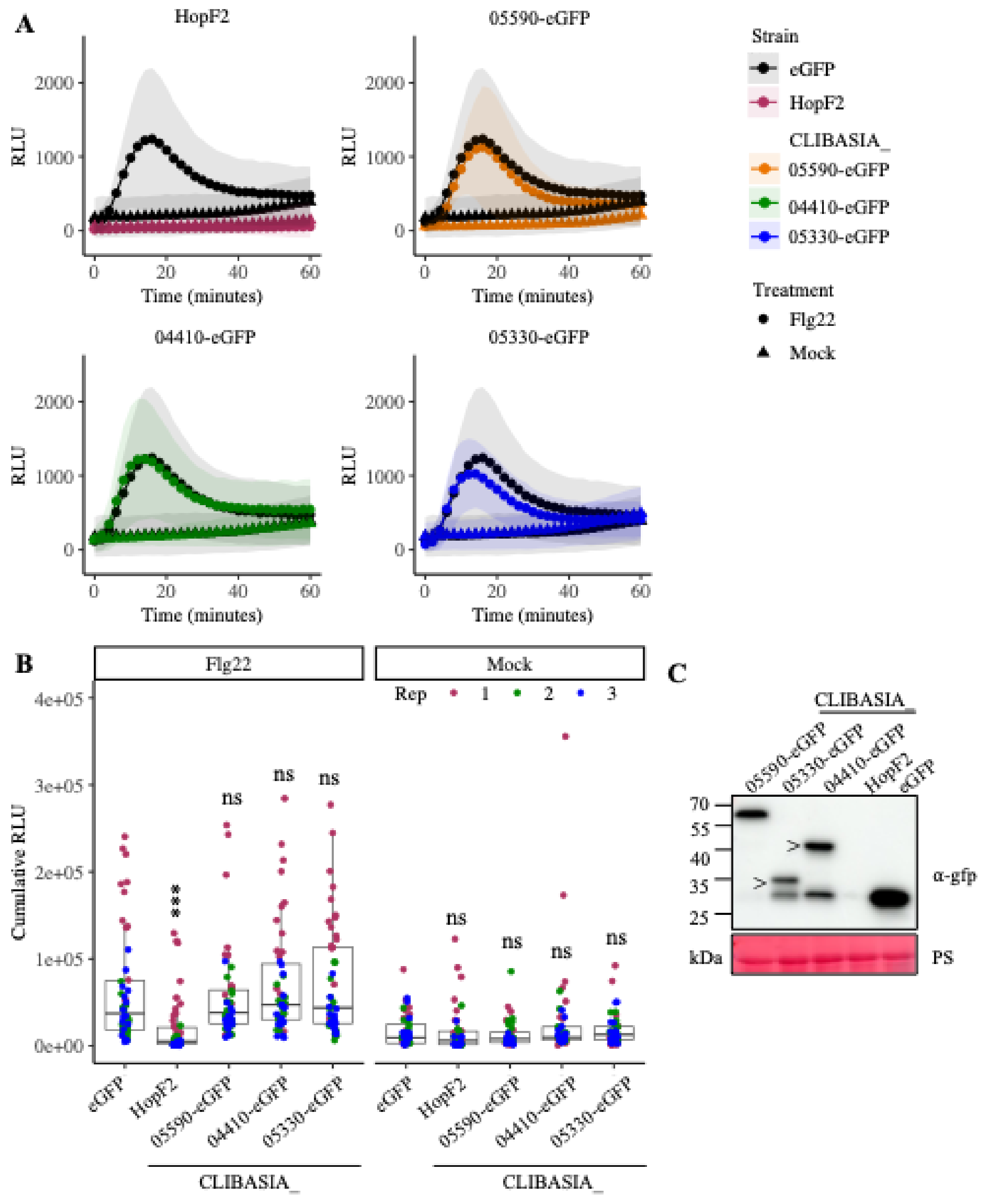


**Figure S2.** Flg22-elicited reactive oxygen species (ROS) burst is unaffected by Las virulence factors when expressed in *Nicotiana benthamiana*. **A-B.** ROS elicited by flg22 was unaffected by Las virulence factors in *N. benthamiana*. Las genes were transiently expressed in *N. benthamiana* leaves through Agroinfiltration. 48 hours post Agroinfiltration, the leaves were treated with 100 nM flg22 (Flg22, circle) or water (Mock, triangle) and total ROS production was evaluated over 60 minutes using a luminol-chemiluminescence assay. eGFP and a *P. syringae* effector HopF2 with known immune suppression activity were used as negative and positive control, respectively. 10 mM MgCl2 was used as the mock treatment.  Shaded regions represent mean ± standard deviation of four disks taken from four plants each. Quantification of the cumulative RLU was determined by measured area under the ROS burst curve. Measurements taken from 4 leaf disks from four plants each and three independent experiments (highlighted by different colored dots). Statistical differences were evaluated based on comparisons to eGFP (2-way ANOVA, ns = not significant, *** $p < 0.001$). **C.** Expression of Las virulence genes in *N. benthamiana* were confirmed by immunoblotting using an anti-GFP antibody. Ponceau Staining (PS) of the membrane was used as a loading control. “>” indicates the correct sized product for samples with multiple bands. HopF2 does not have a tag.

**Supplementary References.**


Baulcombe, D. C., and Molnar, A. 2004. Crystal structure of p19-a universal suppressor of RNA silencing. Trends Biochem. Sci. 29:279-281.

Earley, K. W., Haag, J. R., Pontes, O., Opper, K., Juehne, T., Song, K., and Pikaard, C. S. 2006. Gateway-compatible vectors for plant functional genomics and proteomics. Plant J. 45(4):616-629.

Fleites, L. A., Jain, M., Zhang, S. J. and Gabriel, D. W. 2014. "*Candidatus* Liberibacter asiaticus" prophage late genes may limit host range and culturability. Appl. Environ. Microbiol. 80:6023-6030.

Holsters, M., Silva, B., Van Vliet, F., De Block, M., Dhaese, P., Depicker, A., Inzé, D., Engler, G, Villarroel, R. et al. 1980. The functional organization of the nopaline *A. tumefaciens* plasmid pTiC58. Plasmid. 3:212-230.

Mafra, V., Kubo, K. S., Alves-Ferreira, N., Ribeiro-Alves, M., Stuart, R. M., Boava, L. P., Rodrigues, C. M., and Machado, M. A. 2012. Reference genes for accurate transcript normalization in citrus genotypes under different experimental conditions. PLoS One 7:e31263.

Reyes Caldas, P. A., Zhi, J., Breakspear, A., Thapa, S. P., Toruño, T. Y., Perilla-Henao, L. M., Casteel, C., Faulkner, C. R., and Coaker, G. 2022. Effectors from a bacteria vector-borne pathogen exhibit diverse subcellular localization, expression profiles, and manipulation of plant defense. Mol. Plant-Microbe Interact. 35(12):1067-1080.

Thapa, S. P., De Francesco, A., Trinh, J., Gurung, F. B., Pang, Z., Vidalakis, G., Wang, N., Ancona, V., Ma, W. and Coaker, G. 2020. Genome-wide analyses of Liberibacter species provide insights into evolution, phylogenetic relationships, and virulence factors. Mol. Plant Pathol. 21:716-731.

Yan, Q., Shreedharan, A., Wei, S., Wang, J., Pelz-Stelinski, K., Folimonova, S., and Wang, N. 2013. Global gene expression changes in *Candidatus* Liberibacter asiaticus during the transmission in distinct hosts between plant and insect. Mol. Plant Pathol. 14:391-404.

Yuan, J. and He, S. Y. 1996. The Pseudomonas syringae Hrp regulation and secretion system controls the production and secretion of multiple extracellular proteins. J Bacteriol. 178(21):6399-6402.
